# A comparison of intracranial volume estimation methods and their cross-sectional and longitudinal associations with age

**DOI:** 10.1101/2022.03.29.486254

**Authors:** Stener Nerland, Therese S. Stokkan, Kjetil N. Jørgensen, Laura A. Wortinger, Geneviève Richard, Dani Beck, Dennis van der Meer, Lars T. Westlye, Ole A. Andreassen, Ingrid Agartz, Claudia Barth

**Author notes:** These authors contributed equally (Stener Nerland, Therese S. Stokkan).

## Abstract

Intracranial volume (ICV) is frequently used in volumetric brain magnetic resonance imaging (MRI) studies, both as an adjustment factor for head size and as a variable of interest. Associations with age have been reported in both longitudinal and cross-sectional studies, but results have varied, potentially due to differences in ICV estimation methods. Here, we compared five commonly used ICV estimation methods and their cross-sectional and longitudinal associations with age. T1-weighted cross-sectional MRI data was included for 651 healthy individuals recruited through the NORMENT Centre (mean age = 46.1 years, range = 12.0-85.8 years) and 2,410 healthy individuals recruited through the UK Biobank study (UKB, mean age = 63.2 years, range = 47.0-80.3 years), where follow-up data was also available with a mean follow-up interval of 2.3 years. ICV was estimated with FreeSurfer (eTIV and sbTIV), SPM12, CAT12, and FSL. We assessed Pearson correlations, performed Bland-Altman analysis, and tested the explained variance of sex, height, body weight, and age on pairwise differences between ICV estimation methods. We fitted regression models to test linear and non-linear cross-sectional associations between age and ICV. For the UKB dataset, we further assessed longitudinal ICV change using linear mixed-effects (LME) models. We found overall high correlations across ICV estimation method, with the lowest correlations between FSL and eTIV (r=0.87) and between FSL and CAT12 (r=0.89). Widespread proportional bias was found in the Bland-Altman analyses, i.e., agreement between methods varying as a function of head size. Body weight, age, and sex explained the most variance in the differences between ICV estimation methods, indicating possible confounding by these variables for some estimation methods. In the NORMENT dataset, cross-sectional associations with age were found only for FSL and SPM12, indicating a *positive* association. For the UKB dataset, we observed *negative* cross-sectional associations with age for all ICV estimation methods. Longitudinal associations with age were found for all ICV estimation methods, with estimated annual percentage change ranging from −0.291 % to −0.416 % across the sampled age range. This convergence of longitudinal results across ICV estimation methods, in the largest dataset to date, offers strong evidence for age-related ICV reductions in mid- to late adulthood.

**Highlights:**

- Correlations between the five assessed estimation methods were very high (r>0.90) with the exception of FSL and eTIV (r=0.87), and FSL and CAT12 (r=0.89).
- Explained variance of estimated ICV differences by body weight, age, and sex indicate possible confounding for some ICV estimation methods.
- Positive cross-sectional associations with age, from adolescence to old age, were observed for the SPM12 and FSL estimation methods in one dataset.
- In the other dataset, negative cross-sectional associations with age, from mid- to late adulthood, were found for all estimation methods.
- Longitudinal ICV changes were observed for all estimation methods, indicating an annual percentage ICV reduction of −0.29 % to −0.42 % in mid- to late adulthood.

## 1 Introduction

Intracranial volume (ICV), defined as the volume within the cranium including the brain, meninges and cerebrospinal fluid (CSF), is an important measure in brain magnetic resonance imaging (MRI) studies. It is frequently used to adjust for individual variations in head size (O’Brien et al., 2011; Voevodskaya et al., 2014) and as a proxy for premorbid brain volume in the study of neurodegenerative diseases (Davis & Wright, 1977). Manual delineation of structural brain MRI scans is considered the most accurate *in vivo* method for determining ICV (Huo et al., 2016; Klasson et al., 2015; Whitwell et al., 2001). However, this approach is labor intensive and requires training, making it impractical for large datasets. To overcome these limitations, a variety of automated methods for computing ICV using T1-weighted structural MRI have been developed. Previous studies have reported varying consistency and agreement between ICV estimation methods (Malone et al., 2015; Sargolzaei et al., 2015). In one study, associations between hippocampal volume, education and a cognitive measure differed between ICV estimation methods (Nordenskjöld et al., 2013). Some studies have also indicated that the accuracy of automated ICV estimation varies as a function of head size (Klasson et al., 2018). Such findings highlight the importance of assessing ICV estimation methods for potential sources of bias, which may otherwise introduce spurious effects in studies relying on ICV estimates.

Automated ICV estimation methods typically use T1-weighted MRI images and can be broadly classified as either registration- or segmentation-based. Registration-based methods estimate ICV via an atlas scaling factor given by the determinant of an affine transformation of individual MRI images to a template. The two most common registration-based ICV estimation methods are estimated Total Intracranial Volume (eTIV; Buckner et al., 2004) in FreeSurfer and SIENAX from FSL (FMRIB Software Library; Smith et al., 2002, 2004). With segmentation-based methods, MRI images are first segmented into tissue compartments which are then used to calculate volumetric estimates of the intracranial cavity. One popular segmentation-based method is the Tissue Volumes utility in SPM (Statistical Parametric Mapping; https://www.fil.ion.ucl.ac.uk/spm/). CAT12 is an extension of SPM12 that aims to provide a more robust segmentation algorithm (http://www.neuro.uni-jena.de/cat/). Both SPM12 and CAT12 compute ICV as a probability-weighted sum of grey matter (GM), white matter (WM) and CSF. SAMSEG (Sequence Adaptive Multimodal Segmentation; Puonti et al., 2016) was recently introduced in FreeSurfer version 7 and computes the segmentation-based Total Intracranial Volume (sbTIV).

Throughout the lifespan, ICV increases rapidly from early childhood until early adolescence and is thought to remain relatively stable throughout adulthood (Mills et al., 2016; Pfefferbaum et al., 1994). These findings are consistent with studies on head circumference and computed tomography (Bergerat et al., 2021; Huda et al., 2004; Neubauer et al., 2009). In adulthood, the presence of ICV change is less well-established, and it remains uncertain if continued changes occur and to what extent. However, past non-MRI studies on ICV and head size have often been limited to childhood and adolescence. A notable exception is the cross-sectional study by Weaver and Christian (1980), reporting no significant association between age and occipitofrontal head circumference in 567 participants (50 % female) aged 18 to approximately 67 years.

A number of cross-sectional MRI studies have assessed the association between age and ICV in adulthood, some of which report significant negative associations with age while others find no significant associations. For instance, DeCarli et al. (2005) estimated ICV using in-house software in 2,081 participants from 34 to 96 years of age and found subtle negative associations with age (∼0.1 % ICV change per year), independent of sex. Similarly, in a study of 147 participants between the ages of 15 and 96, Buckner et al (2004) reported a slight negative association between manually determined ICV and age (−1.05 cm^3^ per year). A notable recent example is the study by Ma et al. (2018), where cross-sectional associations with age were analyzed in a large dataset consisting of 7,656 scans for 1,727 elderly subjects (55 to 95 years of age) from the Alzheimer’s Disease Neuroimaging Initiative (ADNI) database. Three diagnostic groups were included; cognitive normal, mild cognitive impairment, and an Alzheimer’s disease group. They observed no significant cross-sectional associations between age and ICV as estimated with three methods: eTIV, SPM12, and multi-atlas label fusion (MALF; Huo et al., 2016).

Most studies on putative associations between ICV and age are cross-sectional rather than longitudinal (Good et al., 2001; Kim et al., 2018; Kruggel, 2006). Such study designs are limited in their ability to resolve age trajectories across the lifespan and can be influenced by generational effects such as secular growth rates (Miller & Corsellis, 1977). In Caspi et al. (2020), both longitudinal and cross-sectional age effects on ICV were assessed in participants between the ages of 16 and 55 years. They included 528 participants at baseline, 378 at follow-up, and 309 at the second follow-up, where the mean period between time points was 3.3 years. In the longitudinal analysis, they reported nonlinear associations where ICV change was initially positive (0.03 % APC at age 20) and then negative in later adulthood (−0.09 % APC at age 55). In the cross-sectional analysis, the data indicated a predicted ICV change of 0.21 % for males and 0.22 % for females at 20 years of age. They attributed the different magnitudes of effects in the longitudinal and cross-sectional analyses to different secular growth rates. We found only one other study on longitudinal ICV changes in the literature, which showed a statistically significant increase in ICV in the youngest group (<= 34 years) but no significant longitudinal age changes in mid- (35-54 years) to late (>= 54 years) adulthood (Liu et al., 2003) where ICV was estimated with Exbrain (Lemieux et al., 2003). However, this study was limited by a small sample size with only 44 participants in the first group, 37 in the second, and 9 in the third.

In the present study, we compared five of the most frequently used automated ICV estimation methods to test their relative consistency and absolute agreement, as well as cross-sectional and longitudinal associations with age. Given the robustness of the ICV measure, we expected that correlation coefficients between estimation methods would exceed 0.9. We further hypothesized that the agreement between ICV estimates would be higher between the two registration-based methods and between the three segmentation-based methods, than across the two groups of ICV estimation methods. To test potential confounding by age, sex, height, and body weight, we computed their explained variance of the pairwise differences between ICV estimation methods. For the cross-sectional associations with age, we expected to find no statistically significant effect of age with segmentation-based ICV estimates and no nonlinear age effects. Based on the previous literature (Buckner et al., 2004; Caspi et al., 2020; DeCarli et al., 2005), we further expected to find negative linear cross-sectional associations with age with a ∼0.15 % decrease in estimated ICV per year with the registration-based methods. Finally, based on the study by Caspi et al., 2020, we expected lower ICV at follow-up compared to baseline with an APC of ∼0.09 %.

## 2 Materials and Methods

### 2.1 Participants and MRI

Participants from two MRI datasets were included in this study: 1) Norwegian Centre for Mental Disorders Research (NORMENT), a cross-sectional dataset recruited in the greater Oslo region of Norway, and 2) UK Biobank (UKB), a longitudinal dataset recruited in the United Kingdom. See **Table 1** for demographics for each dataset and **Fig. 1** for the age distribution for each dataset.

**Fig. 1.**
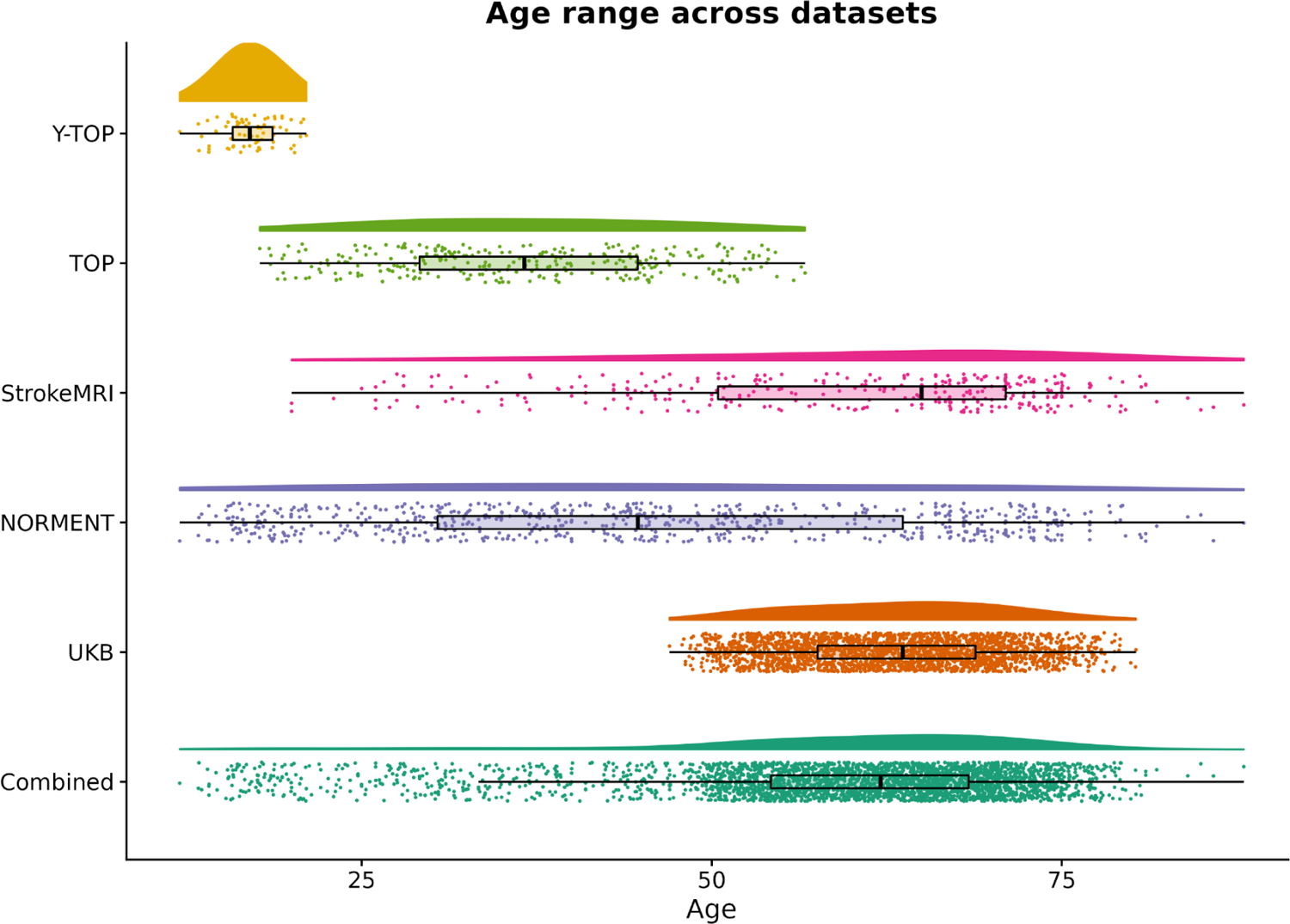
Raincloud plot of the age distributions for each subsample. For UKB we report age at baseline imaging session.

**Table 1.**
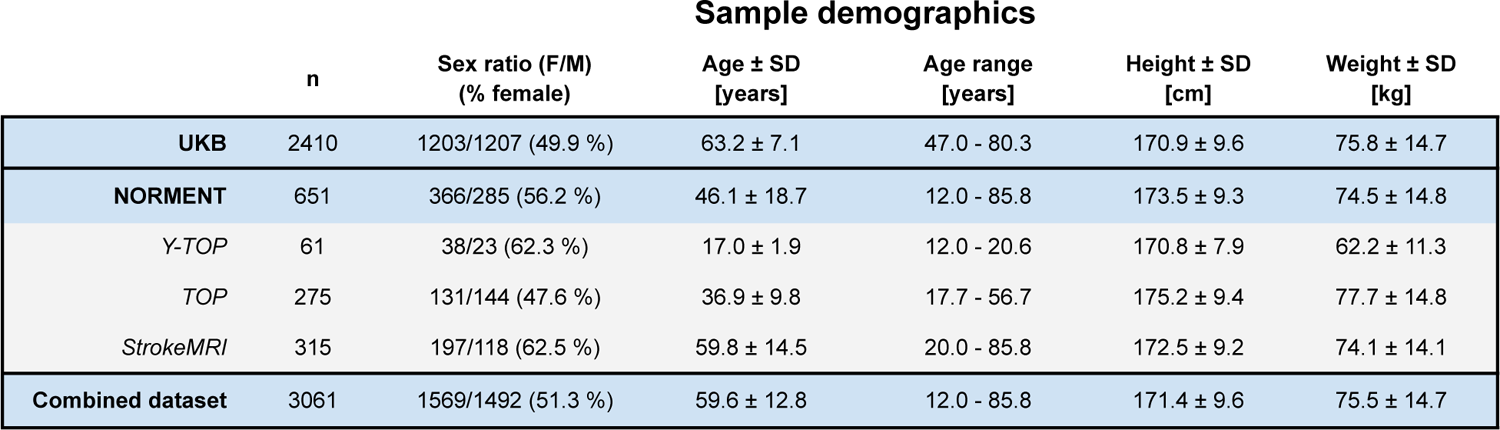
Sample demographics. Continuous variables reported as mean *±* standard deviation. For the UKB dataset, we report age, height, and body weight at baseline imaging session.

#### 2.1.1 NORMENT dataset

Healthy participants were pooled from three clinical studies at the NORMENT Centre: the Youth-Thematically Organized Psychosis (Y-TOP) study (n=61), the Thematically Organized Psychosis (TOP) study (n=275), and the StrokeMRI study (n=315). Participants with complete data, i.e., age, sex, weight, height, and T1-weighted MRI data, were included. For Y-TOP and TOP, invitations to healthy participants were sent out to a random sample, stratified by age and region, from the Norwegian National Population Register in the greater Oslo region. For the StrokeMRI study, healthy participants were recruited via advertisement in local newspapers, social media, and word-of-mouth.

Exclusion criteria for Y-TOP and TOP included a history of neurological disorders or moderate to severe head injury, current or previous diagnosis of a psychiatric disorder, a family history of severe mental disorders, IQ below 70 points, and meeting the criteria for alcohol or substance dependency at the time of MR. For TOP, cannabis use within the last 3 months prior to assessment was an additional exclusion criterion. For StrokeMRI, exclusion criteria included serious head trauma, a history of stroke, dementia or other severe neurological and psychiatric diseases, alcohol- and substance abuse, and medication use thought to affect the nervous system.

Adult participants gave written informed consent to participate. For adolescent participants below 16 years of age, written assent and parental consent was given. Participants were remunerated with a gift card worth 500 NOK. The studies were approved by the Regional Committee for Research Ethics (REK) and the Norwegian Data Protection Authority, and were carried out in accordance with the Helsinki Declaration.

T1-weighted images were acquired at the Oslo University Hospital, Ullevål, on a 3 Tesla General Electric Discovery MR750 scanner, with a 32-channel head coil, between June 2015 and May 2019. An inversion recovery-prepared 3D gradient recalled echo (BRAVO) sequence was employed with the following parameters: repetition time = 8.16 ms, echo time = 3.18 ms, inversion time = 400 ms, field of view = 256 mm, flip angle = 12°, matrix = 188 x 256, voxel size = 1 mm isotropic, 188 sagittal slices.

#### 2.1.2 UKB dataset

Participants with complete longitudinal MRI data and data on age, sex, weight, and height for both baseline and follow-up imaging sessions were selected from the UKB cohort (www.ukbiobank.ac.uk). Time from baseline to follow-up was 2.3 years on average with a standard deviation of 0.1 years. MRI data was acquired at two different sites and participants were scanned at the same site for baseline and follow-up.

We excluded participants with a height difference greater than 5 cm between baseline and follow-up, as we considered this to be indicative of measurement error. Participants were also excluded if they had been diagnosed with disorders known to influence brain structure based on diagnoses from the International Statistical Classification of Diseases and Related Health Problems (ICD-10; World Health Organization, 2004). Diagnostic exclusion criteria included disorders in chapter V and VI, field F; mental and behavioral disorders, including F00 - F03 (Alzheimer’s disease and dementia), F06.7 (mild cognitive disorder), and field G (diseases of the nervous system), including inflammatory and neurodegenerative diseases (except G55-59; ‘Nerve, nerve root and plexus disorders’).

An overview of the UK Biobank acquisition protocols is available in Alfaro-Almagro et al. (2018) and Miller et al. (2016). For MRI, a magnetization prepared rapid acquisition gradient echo (MPRAGE) sequence was employed on a Siemens Skyra 3T scanner with a standard Siemens 32-channel RF receive head coil. The following parameters were used: repetition time = 2000 ms, echo time = 2.01 ms, inversion time = 880 ms, field of view = 256 mm, flip angle = 8°, matrix = 208 x 256, voxel size = 1 mm isotropic.

### 2.2 MRI image processing

T1-weighted MRI images were processed to yield two registration-based ICV measures (eTIV, FSL) and three segmentation-based ICV measures (sbTIV, SPM12 and CAT12). For SPM12 and CAT12, we used MATLAB (The MathWorks, Inc., Massachusetts, USA) version R2018b.

#### 2.2.1 eTIV

We calculated eTIV using the standard processing pipeline, *recon-all*, in FreeSurfer (v5.3.0 for the UKB dataset and v6.0.0 for the NORMENT dataset; https://surfer.nmr.mgh.harvard.edu/). This pipeline performs intensity non-uniformity correction and normalization, skull stripping, and registration to the fsaverage template which is based on the MNI305 template (Evans et al., 1993). The linear scaling factor of this transformation is converted to an ICV estimate by multiplication with the ICV of fsaverage.

#### 2.2.2 FSL

FSL (v6.0.1; https://fsl.fmrib.ox.ac.uk/fsl/) computes ICV with the SIENAX package by first extracting brain and skull images from a single T1-weighted MRI image which is then affinely registered to the MNI152 template (Grabner et al., 2006). Points along the skull are used when determining the registration scaling factor. The final ICV estimate is calculated by multiplying the scaling factor with the measured ICV of the MNI152 template. We ran the *bet* command with a fractional intensity threshold of 0.35 and enabled the bias field and neck cleanup flags.

#### 2.2.3 SPM12

SPM12 (https://www.fil.ion.ucl.ac.uk/spm/) uses a unified segmentation algorithm to perform tissue classification, bias correction and image registration within the same generative model. Based on prior tissue probability maps, it segments the image into tissue classes weighted by the probability of the tissue membership of each tissue type. We used the ‘Tissue Volumes’ utility in SPM12 with the default parameters to calculate ICV as the sum of WM, GM, and CSF.

#### 2.2.4 CAT12

As with SPM12, CAT12 (v12.7; http://www.neuro.uni-jena.de/cat/) uses tissue probability maps to spatially normalize, skull-strip, and initialize the segmentation. In contrast to SPM12, CAT12 uses an adaptive maximum *a posteriori* segmentation approach for determining the final segmentation which accounts for local intensity variations in the original image (Tavares et al., 2020). The goal of this procedure is to provide a segmentation algorithm that is less sensitive to differences in image intensity. As with SPM12, ICV is calculated as the partial volume-adjusted sum of WM, GM and CSF. CAT12 processing was performed with the default parameters.

#### 2.2.5 sbTIV

To compute sbTIV, we used SAMSEG (https://surfer.nmr.mgh.harvard.edu/fswiki/Samseg; Puonti et al., 2016), which creates probability-weighted segmentations of the input image, including skull, non-brain tissue and CSF. sbTIV (https://surfer.nmr.mgh.harvard.edu/fswiki/sbTIV) is computed as a sum of the WM, GM, and CSF volumes.

### 2.3 Statistical analyses

In the combined NORMENT and UKB dataset, we computed Pearson correlations between each ICV estimate and performed Bland-Altman and relative importance analysis. To accommodate for the presence of any sample-specific associations, e.g., due to generational or scanner differences, cross-sectional associations with age were assessed in the two datasets separately. Longitudinal analyses were limited to the UKB dataset. All statistical tests were performed using R Statistical Software (v3.6.3; R Core Team, 2020).

#### 2.3.1 Outlier correction

To avoid excess influence of outliers due to measurement errors, we assessed each ICV estimation method for outliers and excluded the corresponding participants in all subsequent analyses. To identify outliers, we used the median absolute deviation method (Leys et al., 2013) implemented in the R package *Routliers* (https://CRAN.R-project.org/package=Routliers). For the cross-sectional datasets, we used a deviation threshold of 3 on the ICV estimates to identify cross-sectional outliers. For the UKB dataset, we also excluded longitudinal outliers based on the pairwise differences between ICV at baseline and follow-up, where we used a less strict deviation threshold of 4.

We identified and excluded 7 outliers in the NORMENT dataset (1.1 % of sample) and 12 outliers in the cross-sectional UKB dataset (0.5 % of sample). We also identified and excluded 63 longitudinal outliers for the UKB dataset (2.6 % of sample). Two participants were marked both as cross-sectional and longitudinal outliers in the UKB dataset. See **Supplementary Table 1** for demographic information on participants identified as outliers, **Supplementary Figure 1** for an UpSet plot depicting the ICV estimation methods for which outliers were identified, and **Supplementary Note 1** for a description of the outlier detection method and a discussion of the results of outlier correction.

#### 2.3.2 Pearson correlation analyses

To assess pairwise linear relationships between ICV estimation methods in the combined dataset, we calculated Pearson correlation coefficients (r) with 95 % confidence intervals (CI) for each pair of estimation methods using the function *cor.test* from the *stats* R package. To compare the correlations between registration- and segmentation-based methods, we calculated pooled correlations by first applying the Fisher transformation to the correlation coefficients before averaging and back-transforming using the inverse Fisher transformation. We calculated the pooled correlations for each registration-based method with respect to all the segmentation-based methods and compared these pooled correlations to the correlation between the registration-based methods.

#### 2.3.3 Bland-Altman analysis

For each pair of ICV estimation methods in the combined dataset, we created Bland-Altman plots by plotting percentage differences (ΔICV) between ICV method pairs against their means, which can be seen as a proxy for head size. We used 95 % agreement intervals with upper and lower limits of agreement calculated as ΔICV ± 1.96 standard deviation (SD) (Altman & Bland, 1983; Giavarina, 2015). A deviation of ΔICV from zero shows the presence, magnitude, and direction of the difference, or bias, between methods. The bias can be constant or proportional in relation to mean ICV. In the latter case, the difference between methods varies as a function of mean ICV. We quantified these associations by calculating Pearson correlation coefficients between ΔICV and mean ICV and testing their statistical significance.

#### 2.3.4 Relative importance analysis

To assess the influence of age, sex, height, and body weight on the differences between ICV estimates in the combined dataset, we performed relative importance analyses with the *relaimpo* package in R (https://CRAN.R-project.org/package=relaimpo) using the Lindeman, Merenda, and Gold (LMG) metric. This method performs an averaging over the orderings of the explanatory variables to decompose the explained variance of the full model, ICV ∼ Age + Sex + Weight + Height, into the non-negative contributions of each of the variables. The decomposition is constrained to sum to the R^2^ of the full model. The advantage of this method over sequential or nested approaches is that the internal correlational structure of the regressors is taken into account, and the resulting variance decomposition is unbiased (Lindeman et al., 1980).

If neither measure is confounded by age, sex, height, or body weight, we would expect them to explain a negligible proportion of the non-shared variation between them, i.e., the variation of the difference. Thus, we interpreted the degree to which ICV estimate differences were explained by these variables as an indication of possible bias. Note that wherever these variables explain a large proportion of the variance in the difference between two methods, it is not possible to say which of the methods are confounded.

#### 2.3.5 Cross-sectional associations between age and ICV

To test for cross-sectional associations between age and ICV, we first fitted linear regression models for each method with ICV as the outcome variable and age and sex as independent variables. These analyses were conducted separately for the NORMENT and UKB datasets. In additional models, we also included age^2^ to account for possible quadratic associations with age. To visualize the relationship between age and ICV, we created partial regression plots (Velleman & Welsch, 1981) where both age and ICV were residualized with respect to sex and plotted against each other. Cross-sectional Annual Percentage Change (CS-APC) was calculated by expressing the estimated coefficient of the age term in the fitted model as a percentage of the estimated ICV means.

In addition to the linear models, we tested for nonlinear relationships between ICV and age using a generalized additive model (GAM). This method models possible nonlinear relationships using smooth functions (Hastie & Tibshirani, 1986) which can account for higher order nonlinearities. We used cubic regression splines to model age while adjusting for sex. The restricted maximum likelihood method was used as in Sørensen et al. (2021) with the R package *mgcv* (https://CRAN.R-project.org/package=mgcv).

To test the influence of height and body weight on cross-sectional age associations, we fitted additional models including age, sex, height, and body weight as independent variables, as well as separate models with age-by-height and age-by-body weight interactions. To test the influence of sex on age associations, we included models with age, sex, and an age-by-sex interaction term.

To compare the relative qualities of the linear, quadratic and regression spline models, we used the Akaike Information Criterion (AIC; Akaike, 1974). We calculated and compared AIC scores within each ICV estimation method. A low score indicates less information loss in the model, which is a trade-off between goodness of fit and the simplicity of the model. A difference in AIC scores greater than 2 is considered to indicate a significantly better relative quality for the model with the lower score.

Since the age range differed between the NORMENT and the UKB datasets, we performed additional age-matched analyses where we only included a subset of 289 participants in the NORMENT dataset with an age above 47 years. See **Supplementary Tables 7 and 8** for the results of these analyses.

In post-hoc analyses, we ran additional linear regression models where we examined if the age effect differed in the two datasets. Here, ICV was used as the outcome variable and age, sex, and the age-by-cohort interaction as independent variables. The analyses were run both with the complete NORMENT dataset, as well as with the age-matched subset of the NORMENT dataset as described above. The results are reported in **Supplementary Tables 9 and 10**.

#### 2.3.6 Longitudinal associations between age and ICV

To test for longitudinal associations between age and ICV, we fitted linear mixed-effects (LME) models for each ICV estimation method. We entered interscan interval, baseline age, baseline age^2^, scanner, and sex as fixed effects. The primary variable of interest was time point, which provides the contrast between baseline and follow-up. We used individual intercepts as a random effect, in order to allow for between-participant variation in ICV estimates. Annual Percentage Change (APC) was estimated by taking the estimated coefficient of the time point term in the fitted model and expressing it as an annualized percentage of mean ICV.

We fitted additional models to examine the influence of sex, height, and body weight on longitudinal age associations, including a sex-by-time point, height-by-time point, or weight-by-time point interaction term in addition to the terms above. To test the influence of baseline age on longitudinal age associations, we included models with an age-by-time point interaction term.

## 3 Results

### 3.1 Pearson correlation analyses

Pearson correlations between ICV estimation methods were overall high. The lowest correlation coefficient was between eTIV and FSL (r=0.873, CI=[0.865, 0.882]), followed by FSL and CAT12 (r=0.895, CI=[0.887, 0.902]). The highest correlation coefficients were observed between sbTIV and SPM12 (r=0.969, CI=[0.967, 0.971]) and sbTIV and CAT12 (r=0.961, CI=[0.958, 0.963]). See **Fig. 3** for a correlogram showing Pearson correlations with 95 % confidence intervals.

**Fig. 3.**
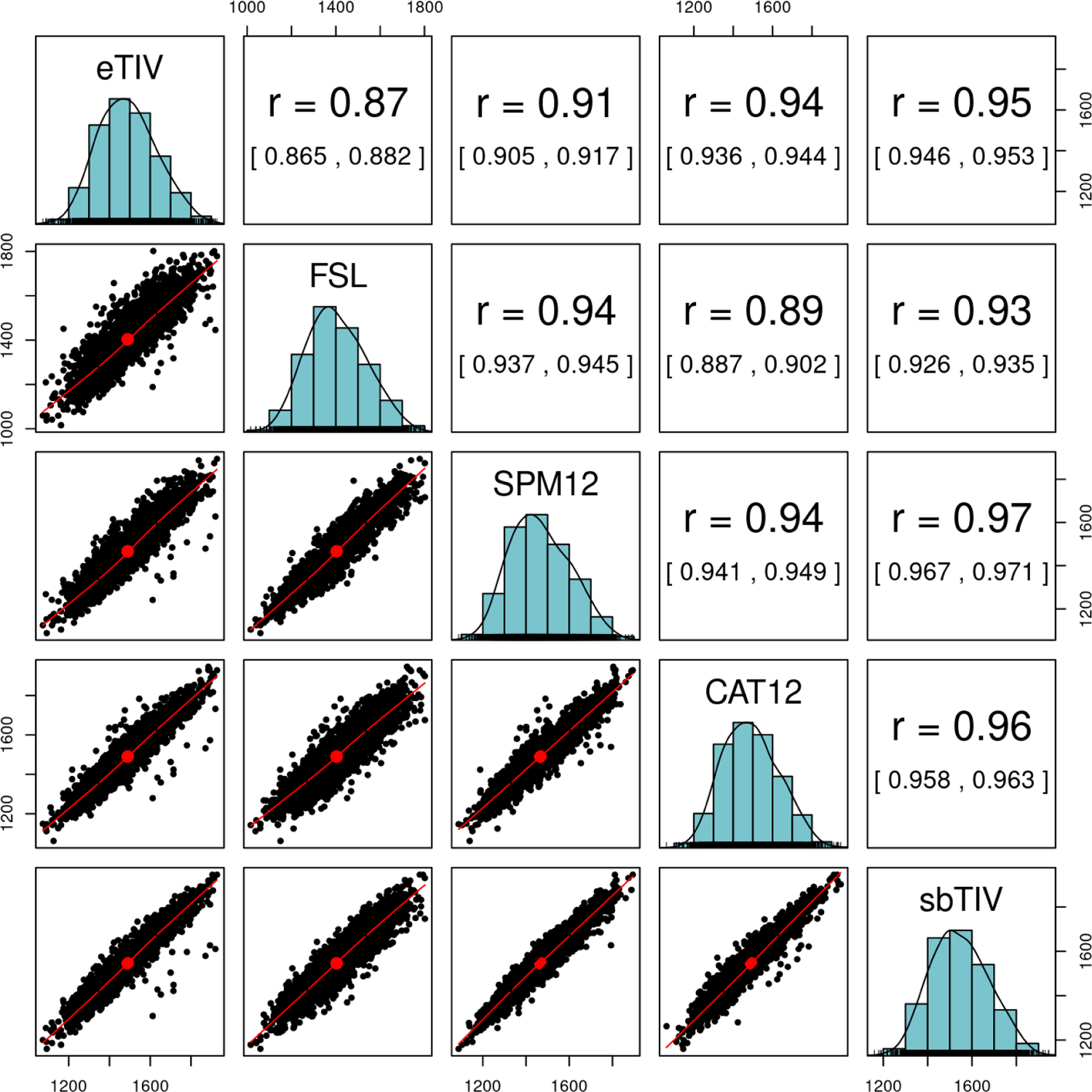
Correlations between each pair of ICV estimation methods in the combined dataset (n=2,981). The main diagonal shows the distribution of each ICV estimate, the lower diagonal shows scatter plots for each pair of ICV estimates, and the top diagonal shows Pearson correlations with 95 % confidence intervals in brackets.

The Pearson correlation between the registration-based methods (eTIV and FSL) was lower (r=0.873) than the pooled correlation between eTIV and the segmentation-based methods (pooled r=0.936) and the pooled correlation between FSL and the segmentation-based methods (pooled r=0.925). The pooled correlation between the segmentation-based methods (SPM12, CAT12, and sbTIV) was 0.959.

### 3.2 Bland-Altman analysis

See **Fig. 4** for Bland-Altman plots. We found that, on average, FSL systematically estimated lower ICV (4.4-9.8 % negative bias) and sbTIV higher ICV (3.8-9.8 % positive bias) compared to the other methods. We also observed statistically significant proportional bias, i.e., correlations between ΔICV and mean ICV for most pairwise comparisons, as indicated by the regression lines in **Fig. 4**. The presence of proportional bias indicates that the agreement between methods differs as a function of the mean ICV across estimation methods. The strongest proportional bias was seen in comparisons of sbTIV with the other ICV estimation methods, where the magnitude of correlations ranged from 0.31 to 0.39 (p<10^-15^). Weaker proportional bias was seen for SPM12 compared to the other methods (excluding sbTIV), here the correlations between ΔICV and mean ICV ranged from 0.05 to 0.04 (p<0.05). We also found weak proportional bias for CAT12 compared to eTIV (r=-0.05, p<0.01).

**Fig. 4.**
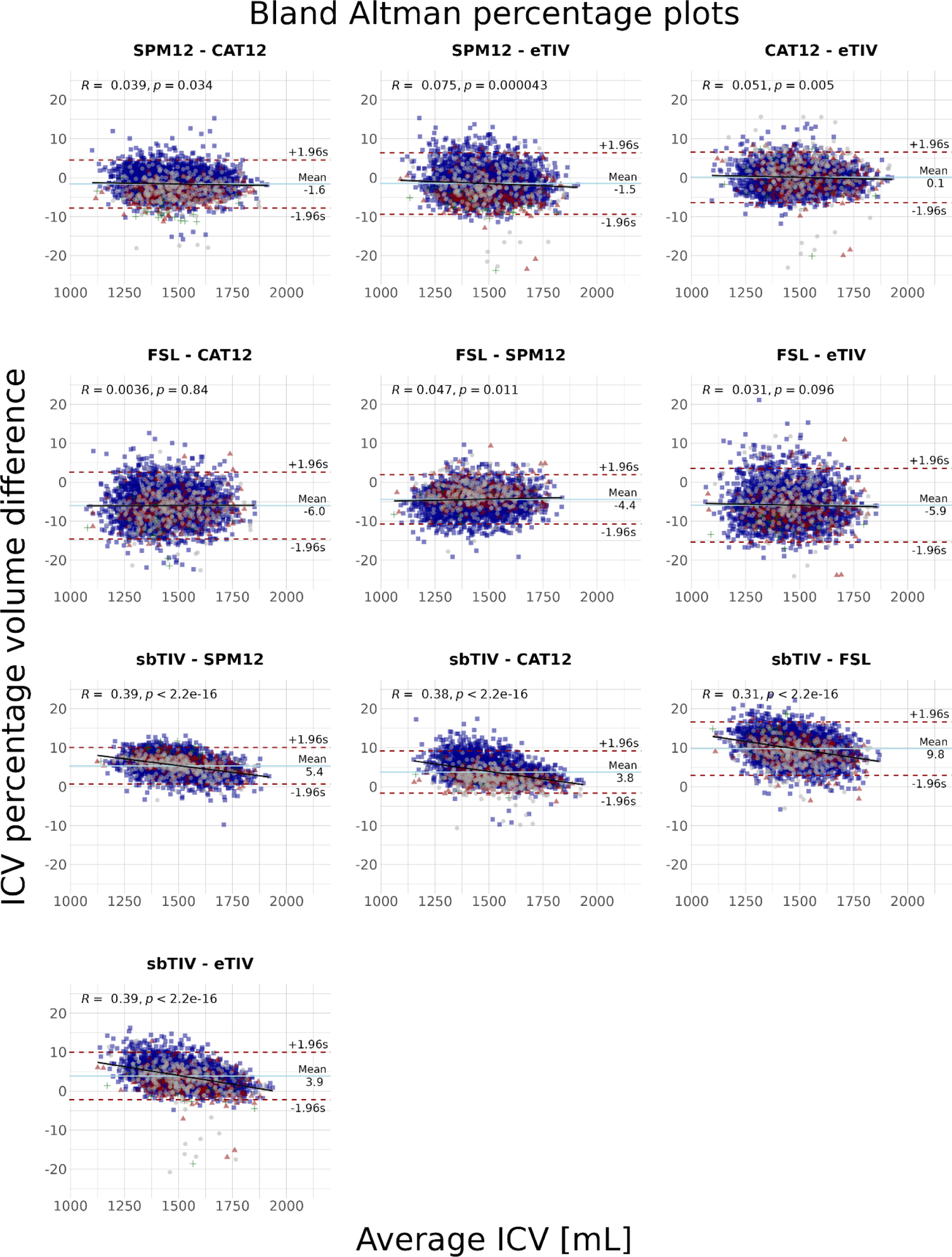
Bland-Altman plots for each pair of ICV estimates in the combined dataset (n=2,981). The means of each pair of estimates (x-axis) are plotted against the percentage differences of the estimates, ΔICV (y-axis). The Pearson correlation coefficient between mean ICV and ΔICV is shown on the top of each plot along with its p-value.

### 3.3 Relative importance analysis

When including age, sex, height, and body weight as explanatory variables, the explained variance of the total model ranged from 2.40 % for the eTIV-CAT12 difference to 28.19 % for the SPM12-sbTIV difference. For the SPM12-sbTIV difference, a large proportion of explained variance was due to sex (9.51 %) and body weight (10.17 %). For the SPM12-CAT12 difference, a large proportion of explained variance was due to body weight (11.36 %) and age (9.08 %). Across pairwise differences, body weight, age, and sex were the best explanatory variables for the differences in ICV estimates. Height explained a small proportion of the variance in most ICV differences, except for the SPM12-sbTIV difference (4.71 %). See **Fig. 5** for bar plots showing the variance decomposition for each explanatory variable.

**Fig. 5.**
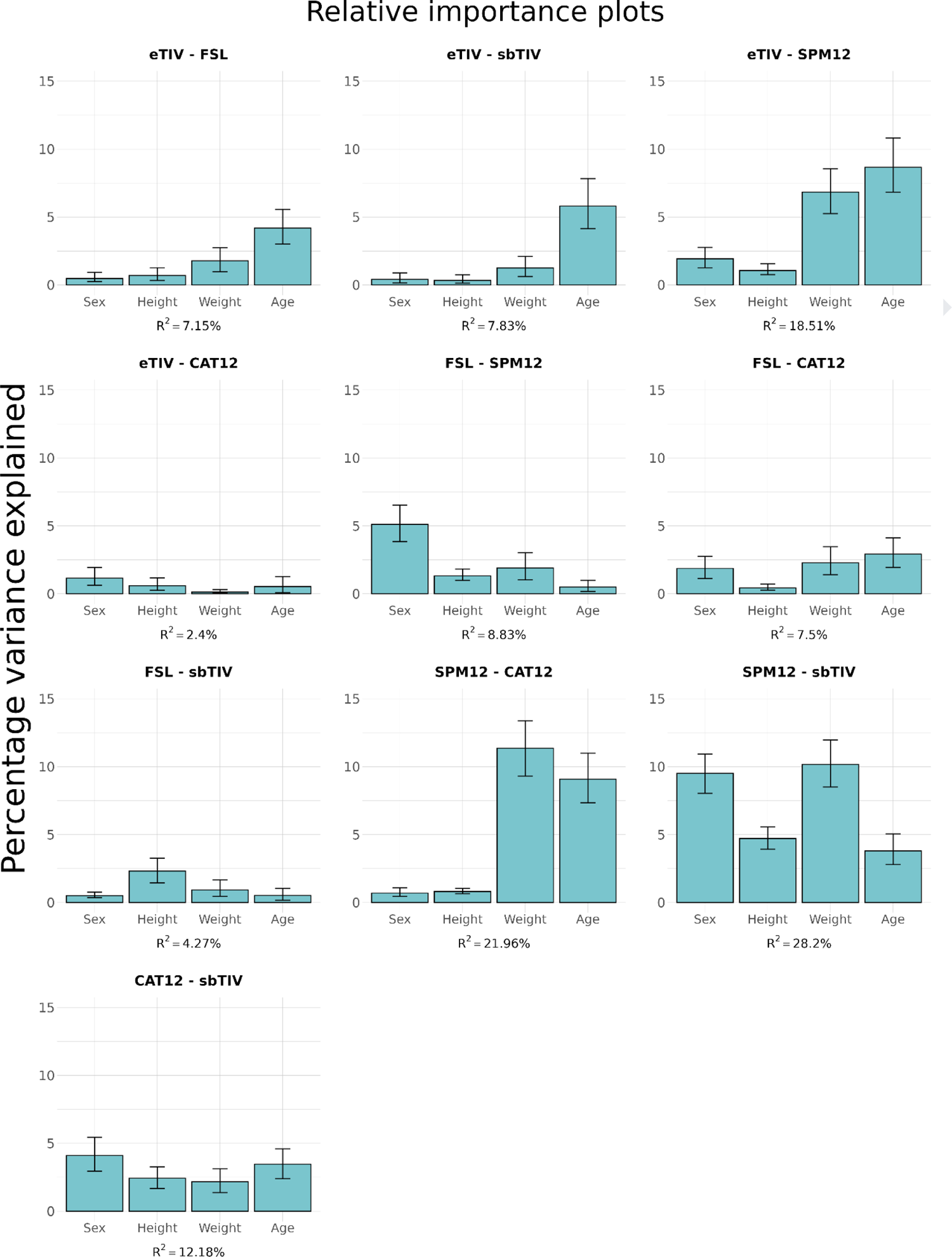
Relative importance of sex, height, body weight, and age on the difference between each ICV measure in the combined dataset (n=2,981).

### 3.4 Cross-sectional associations between age and ICV

Linear models provided the best fit for both datasets and all ICV estimates, except for sbTIV in the UKB dataset where the age^2^ term was significant (p<0.05) and the difference in AIC of the two models exceeded −2, i.e., markedly lower for the quadratic model. See **Supplementary Table 2** for the AIC scores for each model and ICV estimation method for both the NORMENT and the UKB datasets.

For the NORMENT dataset, we found significant *positive* cross-sectional associations with age (ranging from 12.0 to 85.8 years) for FSL (b=1.22, p<10^-6^) and SPM12 (b=1.01, p<10^-5^). These changes corresponded to a CS-APC of 0.086 % for FSL and 0.069 % for SPM12. Age was not significantly associated with any other ICV measure in the NORMENT dataset. For the UKB dataset, we found significant *negative* cross-sectional associations with age (ranging from 47.0 to 80.3 years) for all ICV estimation methods. The greatest CS-APC were seen for CAT12 (−0.107 % CS-APC) and eTIV (−0.107 % CS-APC). The smallest effect size was seen for FSL with an estimated −0.049 % CS-APC.

In the full model, including body weight and height as independent variables in addition to sex and age, we found significant contributions of height for all methods. In the NORMENT dataset, body weight was also significantly associated with ICV for FSL (b=1.27, p<0.001), SPM12 (b=1.38, p<10^-4^), and sbTIV (b=0.87, p<0.05). In the UKB dataset, body weight was associated with ICV for FSL (b=0.51, p<0.01) and SPM12 (b=1.04, p<10^-9^), but not for eTIV, CAT12, or sbTIV. In the NORMENT dataset, the significant associations between age and ICV remained after additionally adjusting for body weight and height for FSL (b=1.06, p<10^-5^) and SPM12 (b=0.84, p<10^-3^). However, in the UKB dataset the cross-sectional associations with age were no longer significant for FSL, SPM12, and sbTIV when adjusting for body weight and height.

In the NORMENT dataset, we observed *positive* interactions between age and sex for SPM12 (b=0.98, p<=0.05) and CAT12 (b=1.07, p<0.05), indicating a more positive slope in males. In the UKB dataset, age-by-sex interactions were found only for FSL (b=-1.52, p<0.05), and in contrast to the findings in the NORMENT dataset, this showed a *negative* interaction between age and sex, indicating a more negative slope in males. We found no significant age-by-height interactions for any ICV estimation method. A significant age-by-weight association was observed for eTIV in the UKB dataset (b=0.05, p<0.05), but not for any of the other ICV estimation methods or in the NORMENT dataset.

In the analyses including a subset of 289 participants from the NORMENT dataset that were age-matched with the UKB dataset, we observed no significant effects of age for any of the ICV estimation methods in the linear model covarying for sex only. When we also adjusted for height and weight, we saw a significant effect of age only for SPM12 (b=1.26, p<0.05).

For the direct comparisons of the age effects between the two datasets, we found significant age-by-cohort interactions for each ICV estimation method when using the complete NORMENT dataset. These interactions indicated a more positive effect of age in the NORMENT dataset compared to the UKB dataset. We also saw a main effect of cohort, indicating higher ICV estimates in the NORMENT dataset compared to the UKB dataset for all ICV estimation methods except sbTIV. When we used the age-matched NORMENT dataset, we found similar significant age-by-cohort interactions for SPM12 (b=1.71, p<0.05), CAT12 (b=2.21, p<0.01), and sbTIV (b=1.63, p<0.05).

See **Fig. 6** and **Fig. 7** for partial regression plots for each dataset where age and ICV are residualized with respect to sex and plotted against each other. See **Supplementary Tables 3 and 4** for further details concerning the main linear regression models used to test cross-sectional associations between ICV and age, and **Supplementary Tables 5 and 6** for the full models including body weight and height. See **Supplementary Tables 7 and 8** for the results of the age-matched analyses and **Supplementary Tables 9 and 10** for the results of the direct comparisons of age effects across datasets.

**Fig. 6.**
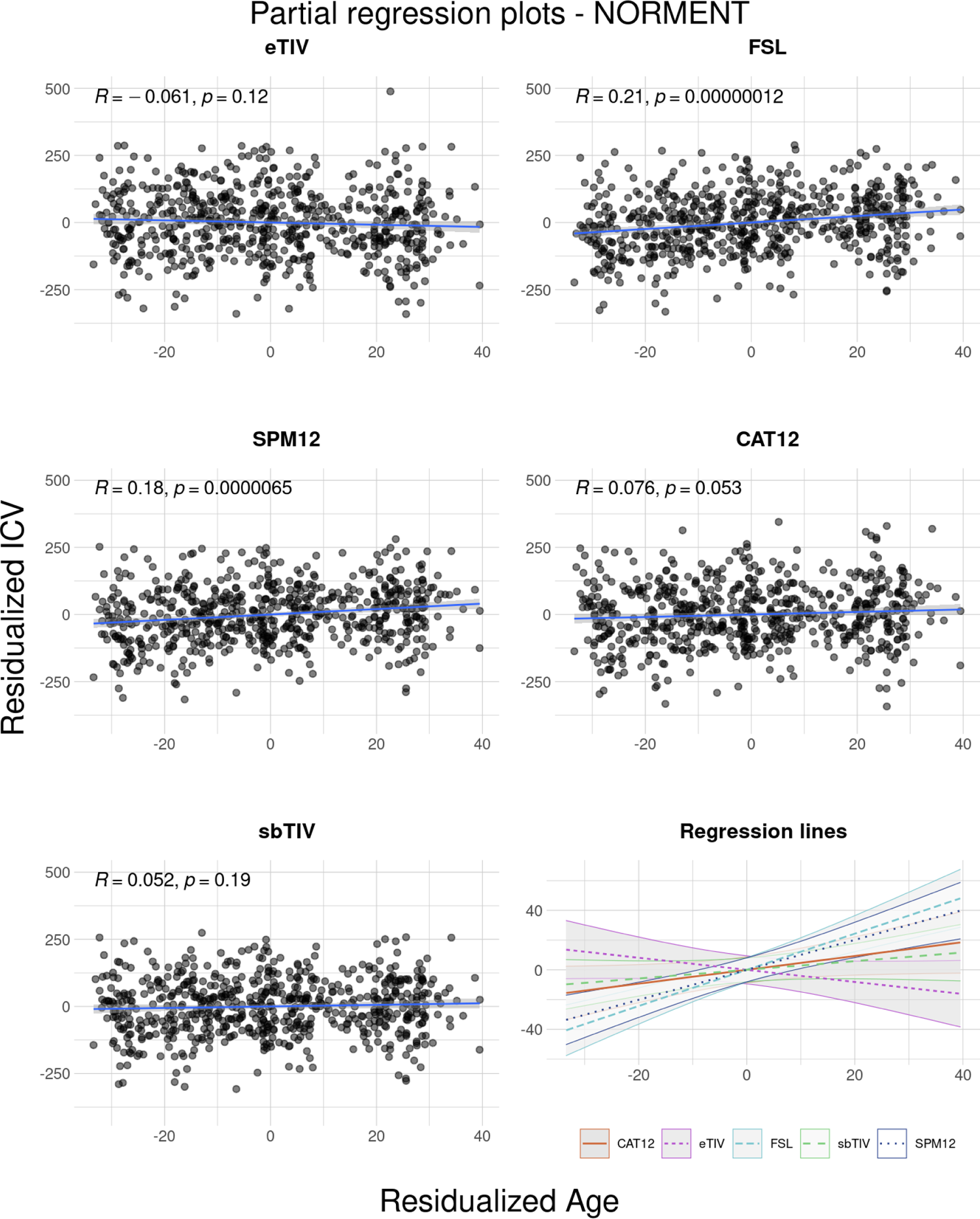
Partial regression plots in the NORMENT dataset (n=644) for the associations between age and ICV for each estimation method. The effect of sex has been regressed out for both age and ICV and the regression lines show the residual effect of age on ICV. As age has been residualised with respect to sex, the x-axis is centered at the sex-adjusted mean.

**Fig. 7.**
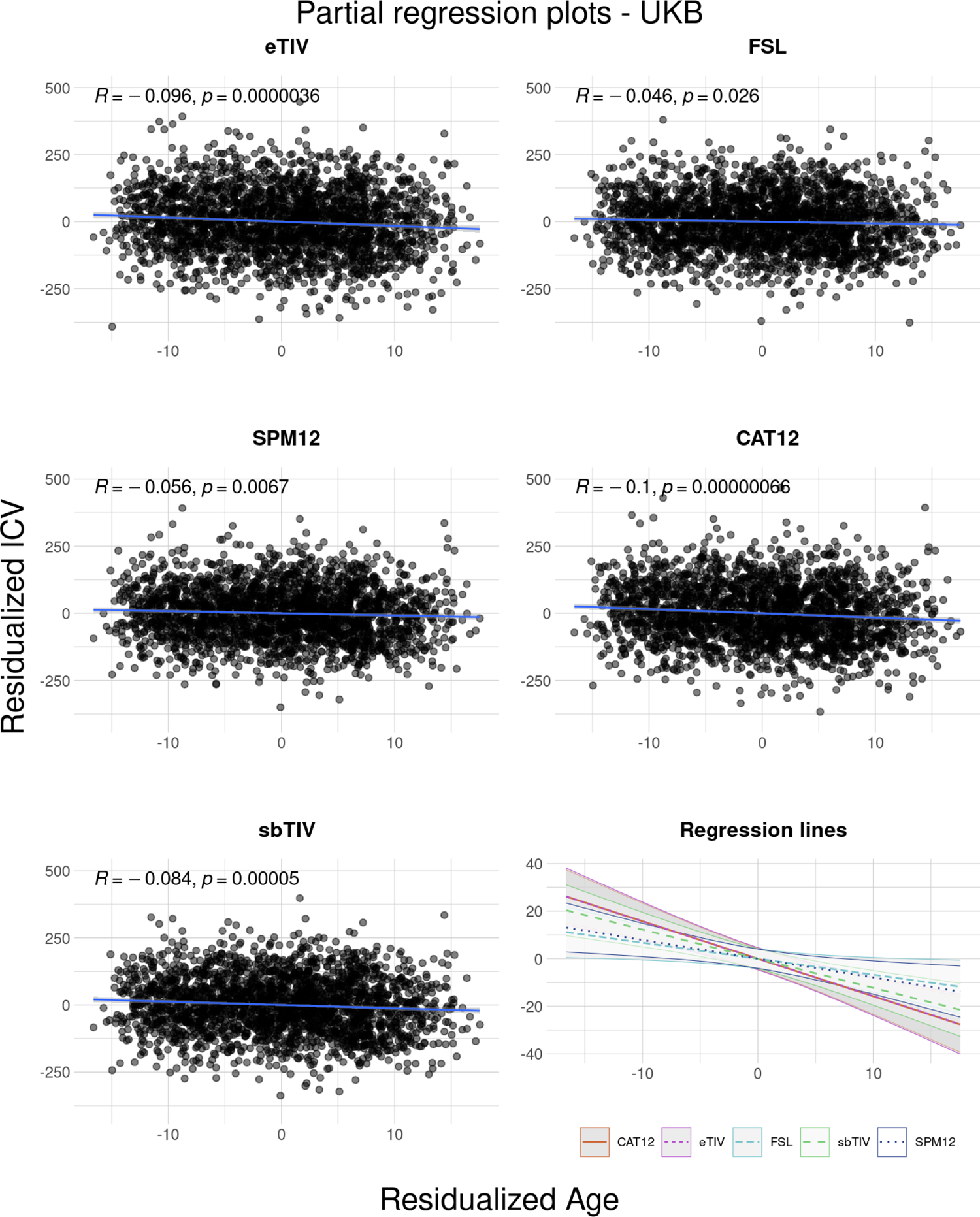
Partial regression plots in the cross-sectional UKB dataset (n=2,337) for the associations between age and ICV for each estimation method. The effect of sex has been regressed out from both age and ICV and the regression lines show the residual effect of age on ICV. As age has been residualised with respect to sex, the x-axis is centered at the sex-adjusted mean.

### 3.5 Longitudinal associations between age and ICV

In the main longitudinal analyses, with fixed factors sex, baseline age, baseline age^2^, scanner, and interscan interval, in the UKB dataset (age range = 47.0-80.3 years), we found significantly lower ICV at follow-up compared to baseline for all ICV estimates. Effect sizes ranged from an APC of −0.291 % for sbTIV to −0.416 % for CAT12. The longitudinal effects of time point on ICV remained for all ICV estimation methods when we included height and weight as fixed factors.

In separate models, we found significant interactions with time point for sex and height for eTIV, but not for any of the other ICV estimation methods. For sex, the interaction indicated lower longitudinal reduction for male participants (b=5.69, p<10^-4^). For height, the interaction showed a lower longitudinal reduction for taller participants (b=0.26, p<0.001). There were no statistically significant interactions between longitudinal ICV change and baseline age.

See **Supplementary Table 1** for details on the main LME model used to investigate longitudinal associations with age. See **Fig. 8** for spaghetti plots depicting longitudinal ICV change in the UKB dataset stratified by sex. See **Supplementary Figure 2** for histograms depicting the raw annual percentage change between baseline and follow-up for each ICV estimation method.

**Fig. 8.**
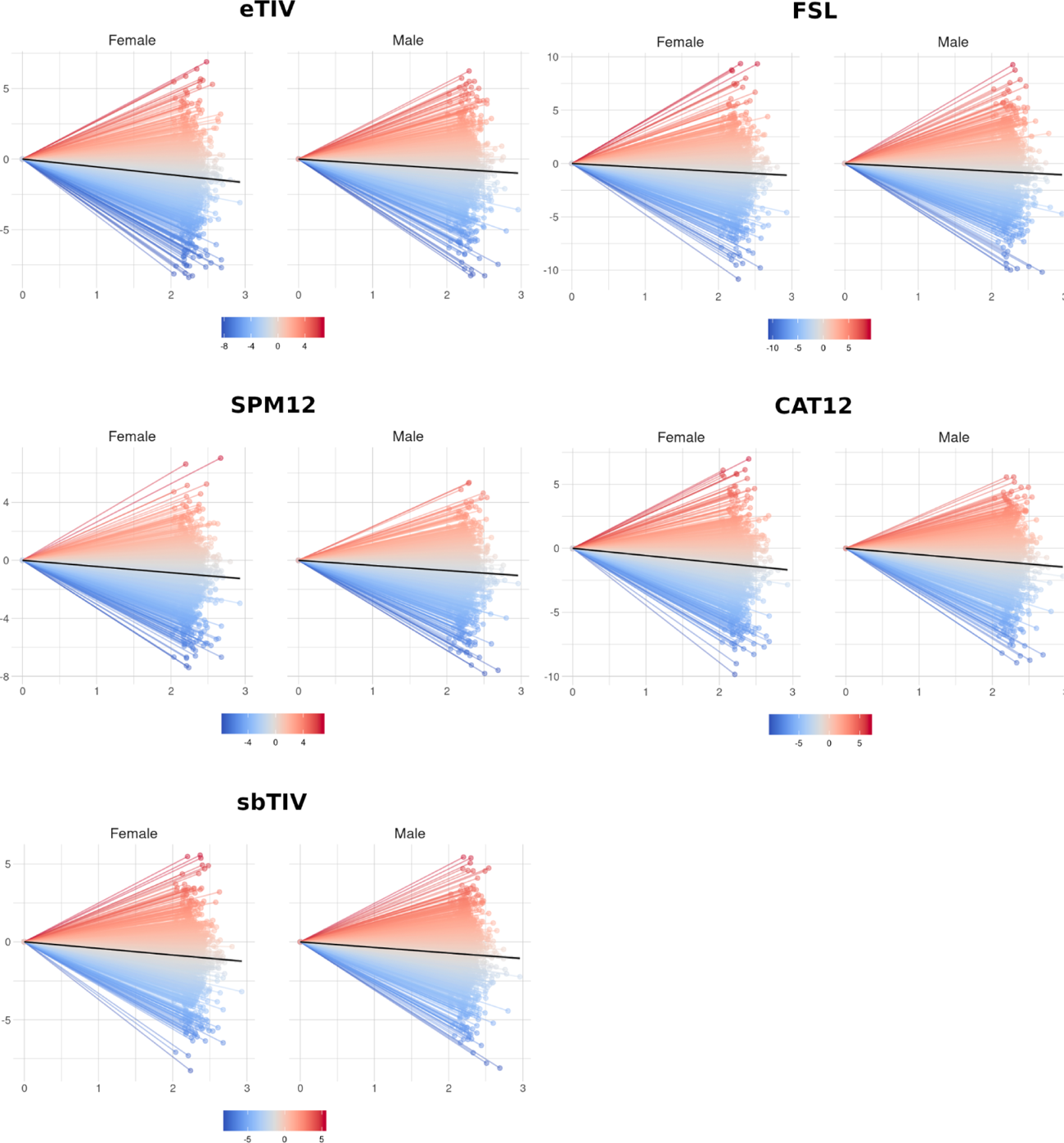
Normalized spaghetti plots depicting longitudinal change for each ICV estimate stratified by sex. The x-axis depicts the interscan interval normalized by time of first imaging session, the y-axis represents ICV change from baseline to follow-up as a percentage of the total ICV across both time points, and the color indicates positive ICV change in red and negative ICV change in blue. The black line depicts the linear trend of longitudinal ICV change.

## 4 Discussion

We compared five commonly used ICV estimation methods, eTIV, FSL, SPM12, CAT12, and SAMSEG, and tested their cross-sectional and longitudinal associations with age. Correlations were overall high, but we found notable differences between the estimation methods. In particular, age, body weight, and sex explained a large proportion for the non-shared variation between several ICV estimation methods. Different cross-sectional effects of age were observed in the two datasets. In the UKB dataset (age range = 47.0-80.3 years), all ICV estimation methods indicated *negative* cross-sectional associations with age, whereas for the NORMENT dataset (age range = 12.0-85.8 years), we observed significant effects only for two of the ICV estimation methods indicating *positive* cross-sectional associations with age. Finally, we observed a striking convergence of results across ICV estimation methods in the longitudinal analyses in the UKB dataset, with an average of 2.3 years from baseline to follow-up, supporting past reports of longitudinal ICV reduction in mid- to late adulthood.

### Correlations between ICV estimation methods

Contrary to our hypothesis, we found higher correlations for each of the registration-based methods, eTIV and FSL, with the segmentation-based methods (eTIV pooled r=0.936; FSL pooled r=0.925) than between eTIV and FSL (r=0.873). The pooled correlations between the segmentation-based methods (SPM12, CAT12, and sbTIV) was 0.959. This points to greater internal consistency within the segmentation-based compared to the registration-based methods. While both eTIV and FSL estimate ICV via an atlas scaling factor, the target template for eTIV is the MNI305-derived fsaverage template (Evans et al., 1993; Fischl et al., 1999), whereas FSL uses the MNI152 template as its target (Mazziotta et al., 1995). Furthermore, FSL uses skull points to constrain the transformation to the template (Smith, 2002; Smith et al., 2001), which may give better registration results compared to eTIV (Buckner et al., 2004). Together, these differences could explain the relatively low correlation between eTIV and FSL, and the differences in their pairwise correlations with the segmentation-based estimation methods.

### Quantitative differences between estimated ICV

Consistent with the previous literature, we found systematic quantitative differences between ICV estimates. In particular, FSL estimated lower ICV, while sbTIV estimated higher ICV than the other methods. Since registration-based methods calculate ICV as the product of an atlas scaling factor and the predetermined ICV of the template, under- or over-estimation of ICV can occur due to the estimated ICV of the template. This could explain the tendency of FSL to underestimate ICV, and a similar overestimation for eTIV relative to manually determined ICV has been reported previously (Klasson et al., 2018). As such, one should be cautious to interpret registration-based methods as absolute measures of ICV.

It is unclear why the segmentation-based method sbTIV appeared to overestimate ICV, but it is worth noting that the anatomical extent of the cranial cavity inferiorly along the spinal cord is not clearly defined, and systematic differences in the cut-off in the different ICV estimation methods may contribute to systematic volumetric differences. Therefore, while segmentation-based methods estimate ICV using a more direct approach, i.e., probability-weighted voxel counts, these methods can also be subject to systematic biases and it remains important to assess their agreement with manually determined ICV if the goal is quantitative ICV estimation.

### Associations between mean ICV and ICV differences

Bland-Altman analyses revealed widespread associations between mean ICV across methods and the differences between them. In other words, the agreement between ICV estimates differed as a function of the mean ICV across methods. Klasson et al. (2018) reported findings consistent with a bias due to head size on eTIV. If mean ICV is considered a proxy for head size, our results lend support to these findings and indicate that similar bias is also present for other ICV estimation methods. It should be noted that while our methods revealed systematic associations in the differences between ICV estimation methods and mean ICV, they cannot say which of the ICV estimation methods have a higher accuracy. We encourage future methodological studies on the validity of automated ICV estimation methods to also include manually determined ICV.

### Relative importance of explanatory variables on ICV differences

In the relative importance analyses, we explored the explained variance of age, sex, body weight and height on the pairwise differences between ICV estimates. If ICV estimation is unbiased by these variables, we would expect to see low explained variance of the estimated ICV differences. Contrary to this expectation, we found a high proportion of explained variance for the total model in the comparison of SPM12 with sbTIV (28.19 %), CAT12 (21.96 %), and eTIV (18.51 %).

Body weight explained 10.17 % of the variance in the SPM12-sbTIV difference and 11.36 % in SPM12-CAT12 difference. This suggests a systematic impact of body weight on ICV differences between these estimation methods. Such bias can be grounds for concern in studies where the variable of interest is associated with weight differences. For example, weight gain is a known side effect of antipsychotic medication (Dayabandara et al., 2017).

We found that sex explained 9.51 % of the SPM12-sbTIV difference. There are contradictory results in the literature on the presence and strength of sexual dimorphism in brain volumes and age-related associations with brain volumes (Fjell et al., 2009; Inano et al., 2013; Ritchie et al., 2018). It remains an open question whether sex-dependent volumetric differences remain after correction for ICV. Past studies have also shown that the statistical correction method can affect sex differences in brain volumes (Pintzka et al., 2015; Sanchis-Segura et al., 2020). The choice of ICV estimation method may explain some of the discrepancy in the past findings. We encourage future studies on sex dimorphism of brain volumes to carefully assess the accuracy of the chosen ICV estimation method, given that confounding by sex can affect results.

### Cross-sectional associations with age

In line with previous studies on cross-sectional age effects on ICV in mid- to late adulthood, we found *negative* cross-sectional associations between ICV and age across all ICV estimation methods in the UKB dataset. Surprisingly, in the NORMENT dataset, which spanned adolescence to late adulthood, we found no statistically significant associations with age for three of the ICV estimation methods (eTIV, CAT12, and sbTIV) and *positive* associations with age for two of the ICV estimation methods (FSL and SPM12). In post hoc analyses, we found statistically significant differences in the age effects in the two datasets for all ICV estimation methods indicating more negative age trajectories in the UKB dataset.

Since the mean age and the age range differed between the UKB and the NORMENT datasets, we performed additional analyses on a subset of 289 participants from the NORMENT dataset that were age-matched to the UKB dataset. In these analyses, none of the main models showed significant associations between age and ICV. However, in the full models, corrected for age, sex, height, and body weight, we saw a significant *positive* association between age and ICV for SPM12 - in line with our findings in the complete dataset. Given the loss of statistical power in these analyses it is difficult to draw firm conclusions. However, the replication of the positive association with age for SPM12 in the full model, and the positive signs of the coefficients for the age term in the main models for FSL and SPM12, suggest that the age associations we observed were not entirely explained by differences in the age ranges. Furthermore, we observed significant age-by-cohort differences in the direct comparisons between the UKB dataset and the age-matched NORMENT dataset for SPM12, CAT12, and sbTIV, but not for eTIV and FSL.

We did not see evidence for non-linear effects of age on ICV in the NORMENT dataset, and the quadratic age term was statistically significant only for sbTIV in the UKB dataset. Past studies on ICV and head size, including computed tomography and head circumference studies (Bergerat et al., 2021; Huda et al., 2004; Neubauer et al., 2009; Weaver and Christian, 1980), lend some support to our findings of linear age trajectories across most ICV estimation methods.

In the NORMENT dataset, we observed sex-by-age interactions for SPM12 and CAT12, indicating a greater cross-sectional ICV increase in males for SPM12. In the UKB dataset, on the other hand, we found a negative sex-by-age interaction for FSL, indicating that male sex is associated with greater cross-sectional ICV decrease. These findings show that sex can, in some cases, affect the cross-sectional associations between age and ICV. This can be a particular concern in clinical group comparisons where sex distributions may vary between groups.

### Longitudinal associations with age

In the longitudinal analyses, we observed statistically significant ICV reduction at follow-up compared to baseline across all estimation methods. Our results indicated an APC ranging from −0.29 % to −0.42 %, which was considerably higher than the APC estimated by Caspi et al. (2020) where they reported an APC of −0.09 % at age 55. However, their age range was from 16 to 55 years of age, and thus only partially overlapped with ours. Notably, they also included people with a diagnosis of schizophrenia and relatives of these participants in the analysis. It is known that ICV has a significant genetic component (Adams et al., 2016), and is affected by genes that confer risk for developing a psychotic disorder (Smeland et al., 2018). Furthermore, cross-sectional studies have found associations between a diagnosis of schizophrenia and ICV (Gurholt et al., 2020; van Erp et al., 2016). Thus, the sample used in Caspi et al. (2020) may be confounded by genetic risk of psychotic disorder, both directly in patients, and indirectly in relatives.

We observed significant sex-by-time point interactions for eTIV, indicating less ICV decrease between baseline and follow-up in males, but no such interactions for the other ICV estimation methods. In isolation, this finding is in line with a semi-longitudinal study of elderly participants (age range=71.1-74.3 years) by Royle et al. (2013) where two sets of manual ICV segmentations were compared in a dataset of elderly participants. The first segmentation measured current ICV, whereas the other was derived from expert manual segmentation where inner table skull thickening (i.e., thickening of the inner bony structure of the skull) was taken into account. The segmentations were used to estimate the longitudinal effect of inner skull thickening on ICV change, yielding an estimated median ICV decrease of 6.2 % in males and 8.3 % in females across the lifespan, in line with the literature on physiological skull changes with age (Harding, 1949). However, we did not observe sex-by-time point interactions with the other ICV estimation methods. It is also unclear whether skull thickening occurs at a constant rate across the lifespan and consequently how much of this effect would be detectable in the age range of the UKB dataset. Furthermore, for eTIV alone we also found a significant height-by-time point interaction indicating less longitudinal change in taller participants. It is possible that the sex-by-time point interaction is driven by height rather than other physiological differences between males and females in this dataset.

We did not find statistically significant baseline age-by-time point interactions for any ICV estimation methods, suggesting that the rate of longitudinal ICV change does not differ with age in mid- to late-adulthood. This finding should, however, be interpreted with caution, given the narrow age range of this sample. In line with the past literature on age-related ICV change, particularly in adolescence, it would be expected that the rate of change differs if longitudinal change is estimated for a greater age range.

### Strengths and limitations

Strengths of our study include the use of large, well-characterized datasets composed of participants with no known neurological or psychiatric disorders thought to affect head size. In particular, the UKB dataset is the largest dataset used for assessing longitudinal ICV change to date. In the cross-sectional analyses, we had a broad age range suitable to assess the presence of nonlinear associations between age and ICV. The data was processed in a harmonized framework, facilitating the direct comparison between ICV estimation methods. For the NORMENT dataset, MRI acquisition was conducted on the same scanner system with no major upgrades.

Some limitations also apply. We processed a total of 5,471 T1-weighted images (651 NORMENT scans + 4,820 UKB baseline and follow-up scans) with five different ICV estimation methods. The resulting dataset of 27,355 ICV estimates precluded detailed quality control of each output. It also prevented us from performing manual segmentation of images, given its time-consuming nature (e.g., ∼30 minutes per subject in Ambarki et al., 2012). Without manually determined ICV estimates, we cannot draw firm conclusions about which estimation methods are more accurate with respect to manual delineation. As with all longitudinal studies, systematic time-dependent bias (e.g., due to scanner drift) can affect the longitudinal results. Finally, our longitudinal dataset only covered a narrow age range, and we did not investigate longitudinal ICV change in adolescence and early adulthood.

## 5 Conclusions

Correlations between ICV estimation methods were lower between the registration-based methods than between segmentation-based methods, suggesting greater internal consistency for the segmentation-based methods. We found that sex, body weight, and age explained a large proportion of the non-shared variation for some pairs of ICV estimation methods. This may represent bias that can be grounds for concern in clinical studies. We observed significant proportional bias for most pairwise comparisons, i.e., varying agreement as a function of mean ICV across methods.

In the NORMENT dataset, spanning adolescence to old age, two ICV estimation methods were *positively* associated with age. In the UKB dataset, spanning mid- to late adulthood, all ICV estimates were *negatively* associated with age. The discovery of age-related effects only for two ICV estimation methods in the NORMENT dataset illustrate how the choice of ICV estimation method can affect findings of age-related associations with ICV. Relationships between ICV and age were significantly different between the two datasets for all estimation methods in the complete dataset, and for three methods in age-matched analyses. This may be due to different secular growth rates in the two cohorts. The convergence of longitudinal results across ICV estimation methods, in the largest dataset to date, offers strong evidence for longitudinal age-related ICV reductions in mid- to late adulthood.

In conclusion, the choice of ICV estimation method is a possible source of bias, not only in studies investigating ICV as a variable of interest, but also in studies that use ICV as an adjustment factor. We encourage future studies to investigate the validity of automated ICV estimation methods and implement quality control procedures to assess the accuracy of the ICV estimation method as with other morphometric brain measures.

## 6 Author contributions

**SN**: Conceptualization, Methodology, Software, Validation, Formal analysis, Investigation, Data curation, Writing - original draft, Writing - review & editing, Visualization, Supervision, Project administration.

**TSS**: Conceptualization, Methodology, Validation, Formal analysis, Investigation, Data curation, Writing - original draft, Writing - review & editing, Visualization.

**KNJ**: Methodology, Writing - original draft, Writing - review & editing.

**LAW**: Writing - original draft, Writing - review & editing.

**GR**: Data curation, Writing - original draft, Writing - review & editing.

**DB**: Writing - original draft, Writing - review & editing.

**DvdM**: Data curation, Writing - original draft, Writing - review & editing.

**LTW**: Resources, Writing - original draft, Writing - review & editing, Funding acquisition.

**OAA**: Resources, Writing - original draft, Writing - review & editing, Funding acquisition.

**IA**: Resources, Writing - original draft, Writing - review & editing, Supervision, Funding acquisition, Project administration.

**CB**: Investigation, Writing - original draft, Writing - review & editing, Supervision, Project administration.

## Acknowledgements

This work was conducted within the Norwegian Centre for Mental Disorders Research, and was funded by the South-Eastern Norway Regional Health Authority; grant number 2017-097, 2019-104, and 2020-020, and the K.G. Jebsen Foundation (grant number SKGJ-MED-008). Access to the UK Biobank dataset was conducted as part of application 27412. Image processing was performed on the TSD (Service for Sensitive Data) computing cluster and at the Imaging Psychosis morphometry lab at Diakonhjemmet Hospital.

## Supplementary Materials

**Supplementary Table 1.** Outlier demographics for each subsample. For the cross-sectional analyses the outliers were identified with outlier detection with the median absolute deviation as implemented in the R package *Routliers* using the standard threshold of 3. For the longitudinal dataset we used a higher threshold of 4 on the ICV differences between baseline and follow-up.

| | n<br>(% sample) | Sex ratio (F/M)<br>(% female) | Age ± SD<br>[years] | Age range<br>[years] | Height ± SD<br>[cm] | Weight ± SD<br>[kg] |
| --- | --- | --- | --- | --- | --- | --- |
| **UKB** | 73 (3.0 %) | 20/53 (27.4 %) | 65.0 ± 7.1 | 49.4 - 78.0 | 176.1 ± 10.5 | 84.2 ± 17.4 |
| **Cross-sectional** | 12 (0.5 %) | 0/12 (0 %) | 65.5 ± 7.3 | 53.9 - 75.5 | 184.3 ± 6.5 | 90.6 ± 13.0 |
| **Longitudinal** | 63 (2.6 %) | 20/43 (31.7 %) | 64.8 ± 7.1 | 49.4 - 78.0 | 174.9 ± 10.6 | 83.4 ± 17.9 |
| **NORMENT** | 7 (1.1 %) | 5/2 (71.4 %) | 58.3 ± 25 | 16.6 - 80.7 | 168.9 ± 6.1 | 71.2 ± 10.6 |

### Supplementary Note 1 - Outlier analysis

To avoid the influence of outliers, we identified and excluded outliers in the statistical analyses using the median absolute deviation method (Leys et al., 2013). Briefly, this method finds the median of the absolute deviations from the median, which yields the median absolute deviation (MAD). Upper and lower thresholds are determined by taking the median of the original dataset ± bᐧMAD where b is the specified threshold (unitless). This approach has the advantage of being robust with respect to sample size and the presence of extreme values.

Since outlier removal can affect results, analyses without outlier correction were also performed. The results of these analyses were similar to those of the outlier-corrected analyses, with a slight decrease in the correlations between ICV estimation methods, especially between eTIV and SPM12. The estimated cross-sectional and longitudinal associations with age were similar for with and without outlier correction.

We found that eTIV was the largest single contributor (i.e., outliers detected for eTIV, but no other ICV estimates) both for cross-sectional outliers in the NORMENT dataset (86 % of outliers) and longitudinal outliers in the UKB dataset (25 % of outliers). In comparison, no participants were marked as outliers for sbTIV alone. This could indicate that eTIV is especially prone to measurement errors and requires a more comprehensive quality control procedure than the other methods, including assessment of the Talairach registration that is performed in the *recon-all* processing stream in FreeSurfer.

**Supplementary Figure 1.**
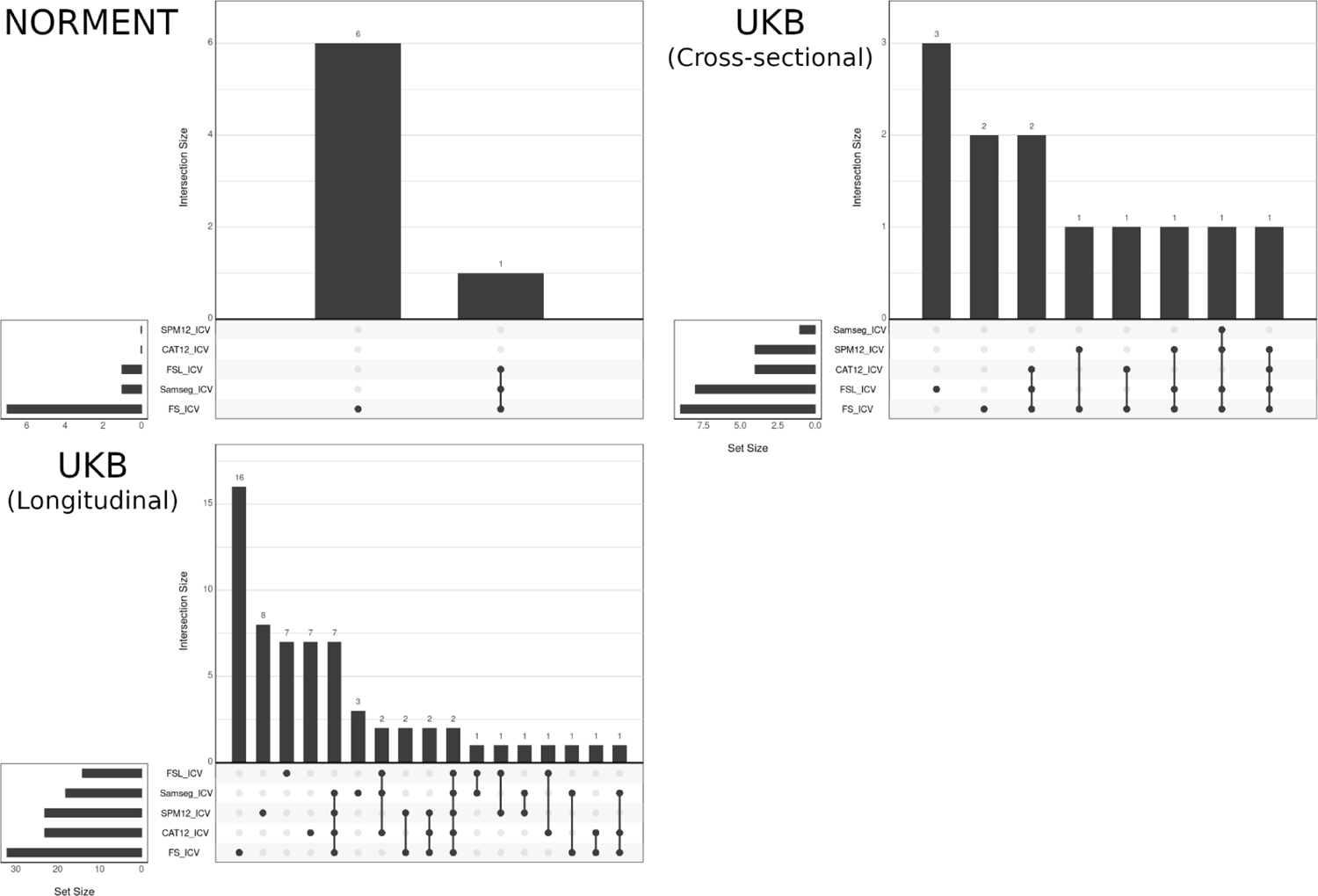
UpSet plots showing the ICV estimation methods for which participants were marked as outliers for the NORMENT dataset, the cross-sectional UKB dataset, and the longitudinal UKB dataset. The height of vertical bars show the number of outliers identified for a particular combination of ICV estimation methods and the length of horizontal bars show the number of times each estimation method was identified as an outlier.

**Supplementary Table 2.** AIC scores and differences with respect to the linear model for each cross-sectional model. A negative difference with magnitude greater than 2 indicates stronger support for the model relative to the linear model. LM = linear model, QM = quadratic model, GAM = generalized additive model, QM - LM = AIC difference between quadratic model and linear model, GAM - LM = AIC difference between generalized additive model and linear model.

| NORMENT |  |  |  |  |  |
| --- | --- | --- | --- | --- | --- |
|  | eTIV | FSL | SPM12 | CAT12 | sbTIV |
| LM | 8,027.03 | 7,850.67 | 7,817.86 | 7,920.65 | 7,816.99 |
| QM | 8,027.26 | 7,850.19 | 7,818.24 | 7,922.61 | 7,818.29 |
| QM - LM | 0.23 | -0.48 | 0.38 | 1.96 | 1.30 |
| GAM | 8,027.35 | 7,850.67 | 7,818.24 | 7,920.66 | 7,817.03 |
| GAM - LM | 0.32 | 0.00 | 0.38 | 0.01 | 0.04 |

**Supplementary Table 2.** AIC scores and differences with respect to the linear model for each cross-sectional model.
| UKB (Cross-sectional) |  |  |  |  |  |
| --- | --- | --- | --- | --- | --- |
|  | eTIV | FSL | SPM12 | CAT12 | sbTIV |
| LM | 28869.71 | 28326.05 | 28145.2 | 28547.7 | 28322.01 |
| QM | 28871.62 | 28325.8 | 28146.07 | 28549.57 | 28319.97 |
| QM - LM | 1.91 | -0.25 | 0.87 | 1.87 | -2.04 |
| GAM | 28879.32 | 28335.91 | 28163.29 | 28559.37 | 28338.55 |
| GAM - LM | 9.61 | 9.86 | 18.09 | 11.67 | 16.54 |

**Supplementary Table 3.** Model details for the main cross-sectional model for each ICV estimation method in the NORMENT dataset. Significant p-values are marked in bold type.

| eTIV |  |  |  |  | FSL |  |  |  |
| --- | --- | --- | --- | --- | --- | --- | --- | --- |
|  | Estimate | Std. error | t-value | p-value | Estimate | Std. error | t-value | p-value |
| Intercept | 1,431.97 | 6.46 | 221.72 | **< 2E-16** | 1,341.90 | 5.63 | 238.27 | **< 2E-16** |
| Sex (M) | 176.09 | 9.74 | 18.08 | **< 2E-16** | 171.73 | 8.49 | 20.22 | **< 2E-16** |
| Age | -0.41 | 0.26 | -1.56 | 0.120 | 1.22 | 0.23 | 5.35 | **1.22E-07** |

| SPM12 |  |  |  |  | CAT12 |  |  |  |
| --- | --- | --- | --- | --- | --- | --- | --- | --- |
|  | Estimate | Std. error | t-value | p-value | Estimate | Std. error | t-value | p-value |
| Intercept | 1,378.48 | 5.49 | 251.08 | **< 2E-16** | 1,438.11 | 5.95 | 241.85 | **< 2E-16** |
| Sex (M) | 190.55 | 8.28 | 23.01 | **< 2E-16** | 178.10 | 8.97 | 19.86 | **< 2E-16** |
| Age | 1.01 | 0.22 | 4.55 | **6.57E-06** | 0.47 | 0.24 | 1.94 | 0.053 |

| sbTIV |  |  |  |  |
| --- | --- | --- | --- | --- |
|  | Estimate | Std. error | t-value | p-value |
| Intercept | 1,467.84 | 5.49 | 267.54 | **< 2E-16** |
| Sex (M) | 176.19 | 8.28 | 21.29 | **< 2E-16** |
| Age | 0.29 | 0.22 | 1.32 | 0.189 |

**Supplementary Table 4.** Model details for the main cross-sectional model for each ICV estimation method in the UKB dataset. Significant p-values are marked in bold type.

| eTIV |  |  |  |  | FSL |  |  |  |
| --- | --- | --- | --- | --- | --- | --- | --- | --- |
|  | Estimate | Std. error | t-value | p-value | Estimate | Std. error | t-value | p-value |
| Intercept | 1,410.99 | 3.87 | 364.90 | < 2E-16 | 1,324.05 | 3.44 | 384.65 | < 2E-16 |
| Sex (M) | 159.27 | 4.82 | 33.02 | < 2E-16 | 164.39 | 4.29 | 38.29 | < 2E-16 |
| Age | -1.58 | 0.34 | -4.69 | **2.95E-06** | -0.68 | 0.30 | -2.27 | **0.023** |
| Scanner | -17.08 | 5.02 | -3.41 | **6.73E-04** | -15.25 | 4.47 | -3.41 | **6.51E-04** |

| SPM12 |  |  |  |  | CAT12 |  |  |  |
| --- | --- | --- | --- | --- | --- | --- | --- | --- |
|  | Estimate | Std. error | t-value | p-value | Estimate | Std. error | t-value | p-value |
| Intercept | 1,382.73 | 3.31 | 417.55 | < 2E-16 | 1,402.49 | 3.61 | 388.57 | < 2E-16 |
| Sex (M) | 186.02 | 4.13 | 45.03 | < 2E-16 | 176.06 | 4.50 | 39.11 | < 2E-16 |
| Age | -0.80 | 0.29 | -2.77 | **0.006** | -1.59 | 0.32 | -5.03 | **5.16E-07** |
| Scanner | -19.10 | 4.30 | -4.45 | **9.19E-06** | -17.31 | 4.68 | -3.70 | **2.24E-04** |

| sbTIV |  |  |  |  |
| --- | --- | --- | --- | --- |
|  | Estimate | Std. error | t-value | p-value |
| Intercept | 1,478.14 | 3.44 | 429.79 | < 2E-16 |
| Sex (M) | 152.46 | 4.29 | 35.54 | < 2E-16 |
| Age | -1.24 | 0.30 | -4.12 | **3.90E-05** |
| Scanner | -19.61 | 4.46 | -4.39 | **1.16E-05** |

**Supplementary Table 5.** Model details for the full cross-sectional model for each ICV estimation method in the NORMENT dataset. Significant p-values are marked in bold type.

| eTIV |  |  |  |  | FSL |  |  |  |
| --- | --- | --- | --- | --- | --- | --- | --- | --- |
|  | Estimate | Std. error | t-value | p-value | Estimate | Std. error | t-value | p-value |
| Intercept | 778.58 | 129.25 | 6.02 | **2.88E-09** | 608.86 | 109.06 | 5.58 | **3.50E-08** |
| Sex (M) | 120.14 | 14.07 | 8.54 | **< 2E-16** | 100.72 | 11.87 | 8.48 | **< 2E-16** |
| Age | -0.45 | 0.26 | -1.74 | 0.083 | 1.06 | 0.22 | 4.85 | **1.56E-06** |
| Weight | 0.28 | 0.40 | 0.71 | 0.480 | 1.27 | 0.34 | 3.75 | **1.95E-04** |
| Height | 3.79 | 0.82 | 4.59 | **5.26E-06** | 3.86 | 0.70 | 5.55 | **4.25E-08** |

| SPM12 |  |  |  |  | CAT12 |  |  |  |
| --- | --- | --- | --- | --- | --- | --- | --- | --- |
|  | Estimate | Std. error | t-value | p-value | Estimate | Std. error | t-value | p-value |
| Intercept | 798.65 | 107.47 | 7.43 | **3.46E-13** | 863.11 | 118.96 | 7.26 | **1.17E-12** |
| Sex (M) | 131.15 | 11.70 | 11.21 | **< 2E-16** | 126.74 | 12.95 | 9.79 | **< 2E-16** |
| Age | 0.84 | 0.22 | 3.91 | **1.03E-04** | 0.40 | 0.24 | 1.67 | 0.095 |
| Weight | 1.38 | 0.34 | 4.13 | **4.14E-05** | 0.50 | 0.37 | 1.34 | 0.181 |
| Height | 2.90 | 0.69 | 4.23 | **2.68E-05** | 3.23 | 0.76 | 4.26 | **2.36E-05** |

| sbTIV |  |  |  |  |
| --- | --- | --- | --- | --- |
|  | Estimate | Std. error | t-value | p-value |
| Intercept | 907.54 | 108.82 | 8.34 | **4.57E-16** |
| Sex (M) | 122.80 | 11.85 | 10.37 | **< 2E-16** |
| Age | 0.18 | 0.22 | 0.84 | 0.400 |
| Weight | 0.87 | 0.34 | 2.57 | **0.010** |
| Height | 2.99 | 0.69 | 4.31 | **1.89E-05** |

**Supplementary Table 6.** Model details for the main cross-sectional model for each ICV estimation method in the cross-sectional UKB dataset. Significant p-values are marked in bold type.

| eTIV |  |  |  |  | FSL |  |  |  |
| --- | --- | --- | --- | --- | --- | --- | --- | --- |
|  | Estimate | Std. error | t-value | p-value | Estimate | Std. error | t-value | p-value |
| Intercept | 804.92 | 62.81 | 12.81 | **< 2E-16** | 496.54 | 54.09 | 9.18 | **< 2E-16** |
| Sex (M) | 107.28 | 7.27 | 14.76 | **< 2E-16** | 87.66 | 6.26 | 14.01 | **< 2E-16** |
| Age | -0.93 | 0.34 | -2.75 | **0.006** | 0.30 | 0.29 | 1.02 | 0.308 |
| Weight | -0.09 | 0.20 | -0.45 | 0.657 | 0.51 | 0.17 | 2.95 | **0.003** |
| Height | 3.73 | 0.40 | 9.28 | **< 2E-16** | 4.83 | 0.35 | 13.95 | **< 2e-16** |
| Scanner | -13.55 | 4.95 | -2.74 | **0.006** | -11.40 | 4.26 | -2.68 | **0.007** |

| SPM12 |  |  |  |  | CAT12 |  |  |  |
| --- | --- | --- | --- | --- | --- | --- | --- | --- |
|  | Estimate | Std. error | t-value | p-value | Estimate | Std. error | t-value | p-value |
| Intercept | 691.35 | 52.24 | 13.24 | **< 2E-16** | 780.23 | 58.39 | 13.36 | **< 2E-16** |
| Sex (M) | 116.38 | 6.04 | 19.26 | **< 2E-16** | 124.93 | 6.76 | 18.49 | **< 2E-16** |
| Age | 0.10 | 0.28 | 0.37 | 0.713 | -0.95 | 0.31 | -3.03 | **0.002** |
| Weight | 1.04 | 0.17 | 6.19 | **7.25E-10** | -0.34 | 0.19 | -1.81 | 0.070 |
| Height | 3.79 | 0.33 | 11.31 | **< 2E-16** | 3.94 | 0.37 | 10.52 | **< 2E-16** |
| Scanner | -16.83 | 4.11 | -4.09 | **4.43E-05** | -13.29 | 4.60 | -2.89 | **0.004** |

| sbTIV |  |  |  |  |
| --- | --- | --- | --- | --- |
|  | Estimate | Std. error | t-value | p-value |
| Intercept | 824.13 | 55.21 | 14.93 | **< 2E-16** |
| Sex (M) | 92.63 | 6.39 | 14.50 | **< 2E-16** |
| Age | -0.48 | 0.30 | -1.60 | 0.109 |
| Weight | 0.32 | 0.18 | 1.78 | 0.075 |
| Height | 3.86 | 0.35 | 10.90 | **< 2E-16** |
| Scanner | -16.43 | 4.35 | -3.78 | **1.62E-04** |

**Supplementary Table 7.**
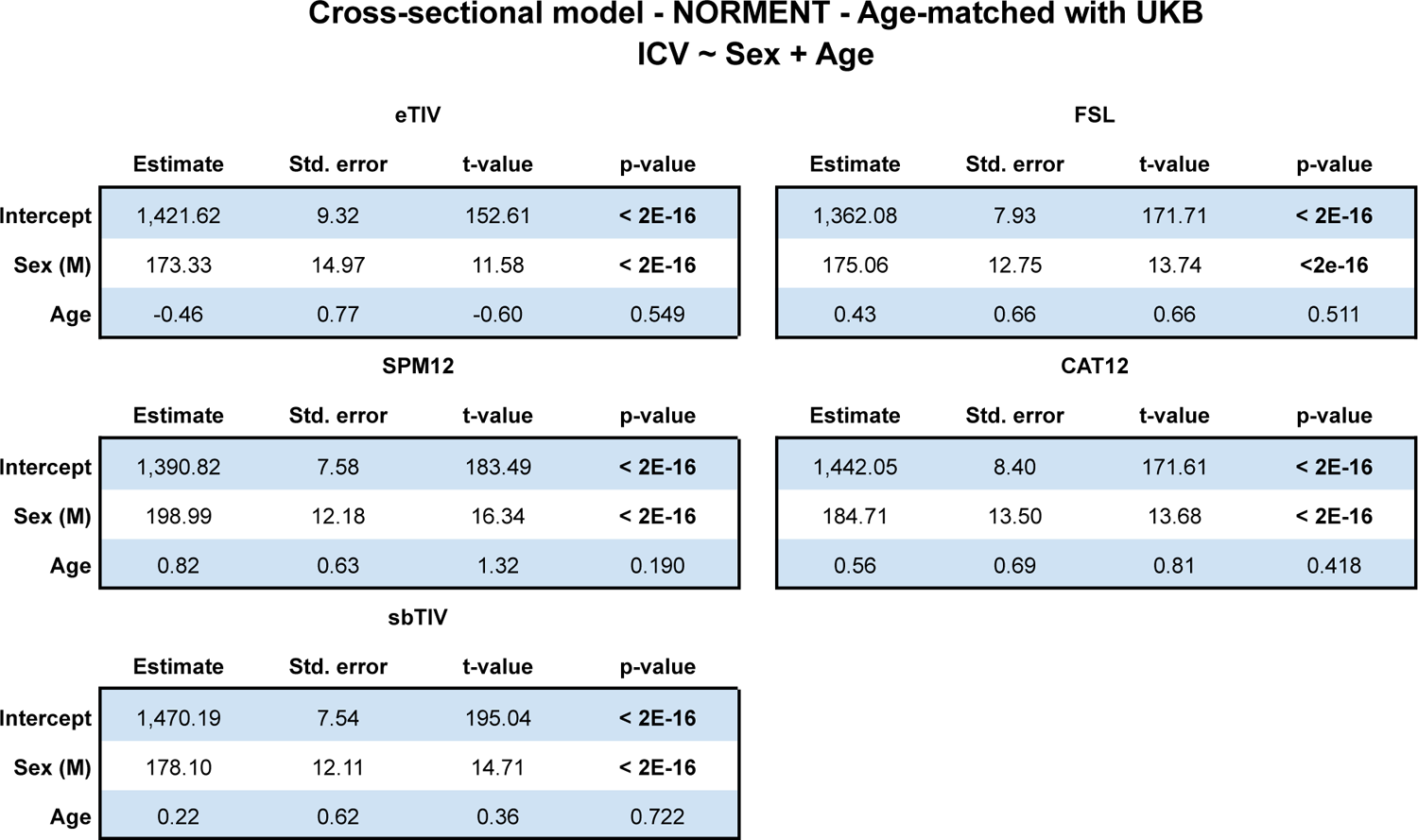
Model details for the main cross-sectional model for each ICV estimation method in the NORMENT dataset age-matched with the UKB dataset. Significant p-values are marked in bold type.

**Cross-sectional model - NORMENT - Age-matched with UKB ICV ~ Sex + Age**
| eTIV |  |  |  |  | FSL |  |  |  |
| --- | --- | --- | --- | --- | --- | --- | --- | --- |
|  | Estimate | Std. error | t-value | p-value | Estimate | Std. error | t-value | p-value |
| Intercept | 1,421.62 | 9.32 | 152.61 | **< 2E-16** | 1,362.08 | 7.93 | 171.71 | **< 2E-16** |
| Sex (M) | 173.33 | 14.97 | 11.58 | **< 2E-16** | 175.06 | 12.75 | 13.74 | **<2e-16** |
| Age | -0.46 | 0.77 | -0.60 | 0.549 | 0.43 | 0.66 | 0.66 | 0.511 |

| SPM12 |  |  |  |  | CAT12 |  |  |  |
| --- | --- | --- | --- | --- | --- | --- | --- | --- |
|  | Estimate | Std. error | t-value | p-value | Estimate | Std. error | t-value | p-value |
| Intercept | 1,390.82 | 7.58 | 183.49 | **< 2E-16** | 1,442.05 | 8.40 | 171.61 | **< 2E-16** |
| Sex (M) | 198.99 | 12.18 | 16.34 | **< 2E-16** | 184.71 | 13.50 | 13.68 | **< 2E-16** |
| Age | 0.82 | 0.63 | 1.32 | 0.190 | 0.56 | 0.69 | 0.81 | 0.418 |

| sbTIV |  |  |  |  |
| --- | --- | --- | --- | --- |
|  | Estimate | Std. error | t-value | p-value |
| Intercept | 1,470.19 | 7.54 | 195.04 | **< 2E-16** |
| Sex (M) | 178.10 | 12.11 | 14.71 | **< 2E-16** |
| Age | 0.22 | 0.62 | 0.36 | 0.722 |

**Supplementary Table 8.** Model details for the full cross-sectional model for each ICV estimation method in the NORMENT dataset age-matched with the UKB dataset. Significant p-values are marked in bold type.

| eTIV |  |  |  |  | FSL |  |  |  |
| --- | --- | --- | --- | --- | --- | --- | --- | --- |
|  | Estimate | Std. error | t-value | p-value | Estimate | Std. error | t-value | p-value |
| Intercept | 1,006.14 | 209.08 | 4.81 | **2.43E-06** | 626.02 | 171.51 | 3.65 | **3.12E-04** |
| Sex (M) | 141.14 | 22.63 | 6.24 | **1.60E-09** | 104.55 | 18.56 | 5.63 | **4.26E-08** |
| Age | -0.29 | 0.78 | -0.38 | 0.708 | 0.95 | 0.64 | 1.49 | 0.138 |
| Weight | -0.23 | 0.70 | -0.34 | 0.738 | 0.95 | 0.57 | 1.66 | 0.098 |
| Height | 2.58 | 1.35 | 1.91 | 0.057 | 4.01 | 1.11 | 3.62 | **3.47E-04** |

| SPM12 |  |  |  |  | CAT12 |  |  |  |
| --- | --- | --- | --- | --- | --- | --- | --- | --- |
|  | Estimate | Std. error | t-value | p-value | Estimate | Std. error | t-value | p-value |
| Intercept | 805.89 | 165.78 | 4.86 | **1.93E-06** | 886.36 | 186.90 | 4.74 | **3.35E-06** |
| Sex (M) | 141.37 | 17.94 | 7.88 | **7.04E-14** | 139.24 | 20.23 | 6.88 | **3.73E-11** |
| Age | 1.26 | 0.62 | 2.04 | **0.042** | 0.83 | 0.70 | 1.19 | 0.234 |
| Weight | 0.92 | 0.55 | 1.66 | 0.099 | -0.07 | 0.62 | -0.11 | 0.912 |
| Height | 3.12 | 1.07 | 2.92 | **3.84E-03** | 3.35 | 1.21 | 2.78 | **0.006** |

| sbTIV |  |  |  |  |
| --- | --- | --- | --- | --- |
|  | Estimate | Std. error | t-value | p-value |
| Intercept | 994.16 | 166.78 | 5.96 | **7.43E-09** |
| Sex (M) | 131.57 | 18.05 | 7.29 | **3.10E-12** |
| Age | 0.57 | 0.62 | 0.92 | 0.358 |
| Weight | 0.71 | 0.56 | 1.27 | 0.205 |
| Height | 2.55 | 1.08 | 2.37 | **0.018** |

**Supplementary Table 9.** Model details for the age-by-cohort interaction with the complete NORMENT dataset. Significant p-values are marked in bold type.

| eTIV |  |  |  |  | FSL |  |  |  |
| --- | --- | --- | --- | --- | --- | --- | --- | --- |
|  | Estimate | Std. error | t-value | p-value | Estimate | Std. error | t-value | p-value |
| Intercept | 1,408.79 | 3.45 | 408.89 | < 2E-16 | 1,320.16 | 3.05 | 432.60 | < 2E-16 |
| Sex (M) | 163.22 | 4.33 | 37.66 | < 2E-16 | 166.28 | 3.84 | 43.32 | < 2E-16 |
| Age | -1.59 | 0.34 | -4.63 | **3.76E-06** | -0.68 | 0.30 | -2.24 | **0.025** |
| Cohort (NORMENT) | 23.09 | 6.38 | 3.62 | **2.98E-04** | 40.92 | 5.65 | 7.24 | **5.52E-13** |
| Age-by-cohort | 1.17 | 0.42 | 2.76 | **0.006** | 1.89 | 0.38 | 5.03 | **5.19E-07** |

| SPM12 |  |  |  |  | CAT12 |  |  |  |
| --- | --- | --- | --- | --- | --- | --- | --- | --- |
|  | Estimate | Std. error | t-value | p-value | Estimate | Std. error | t-value | p-value |
| Intercept | 1,378.13 | 2.95 | 467.41 | < 2E-16 | 1,401.74 | 3.21 | 437.26 | < 2E-16 |
| Sex (M) | 187.39 | 3.71 | 50.52 | < 2E-16 | 176.85 | 4.03 | 43.86 | < 2E-16 |
| Age | -0.79 | 0.29 | -2.70 | **0.007** | -1.58 | 0.32 | -4.95 | **7.72E-07** |
| Cohort (NORMENT) | 15.64 | 5.46 | 2.87 | **0.004** | 43.34 | 5.93 | 7.31 | **3.55E-13** |
| Age-by-cohort | 1.80 | 0.36 | 4.95 | **7.88E-07** | 2.04 | 0.39 | 5.17 | **2.49E-07** |

| sbTIV |  |  |  |  |
| --- | --- | --- | --- | --- |
|  | Estimate | Std. error | t-value | p-value |
| Intercept | 1,473.01 | 3.04 | 484.74 | < 2E-16 |
| Sex (M) | 157.95 | 3.82 | 41.32 | < 2E-16 |
| Age | -1.25 | 0.30 | -4.13 | **3.78E-05** |
| Cohort (NORMENT) | 6.71 | 5.62 | 1.19 | 0.233 |
| Age-by-cohort | 1.52 | 0.37 | 4.07 | **4.75E-05** |

**Supplementary Table 10.** Model details for the age-by-cohort interaction with the age-matched NORMENT dataset. Significant p-values are marked in bold type.

| eTIV |  |  |  |  | FSL |  |  |  |
| --- | --- | --- | --- | --- | --- | --- | --- | --- |
|  | Estimate | Std. error | t-value | p-value | Estimate | Std. error | t-value | p-value |
| Intercept | 1,403.86 | 3.33 | 422.18 | < 2E-16 | 1,317.81 | 2.95 | 447.48 | < 2E-16 |
| Sex (M) | 161.21 | 4.60 | 35.02 | < 2E-16 | 165.93 | 4.08 | 40.70 | < 2E-16 |
| Age | -1.59 | 0.34 | -4.66 | **3.29E-06** | -0.69 | 0.30 | -2.27 | **0.023** |
| Cohort (NORMENT) | 22.69 | 7.33 | 3.09 | **0.002** | 47.63 | 6.49 | 7.33 | **2.97E-13** |
| Age-by-cohort | 1.21 | 0.80 | 1.51 | 0.131 | 1.18 | 0.71 | 1.67 | 0.096 |

| SPM12 |  |  |  |  | CAT12 |  |  |  |
| --- | --- | --- | --- | --- | --- | --- | --- | --- |
|  | Estimate | Std. error | t-value | p-value | Estimate | Std. error | t-value | p-value |
| Intercept | 1,374.94 | 2.84 | 484.94 | < 2E-16 | 1,395.55 | 3.09 | 451.29 | < 2E-16 |
| Sex (M) | 187.92 | 3.93 | 47.87 | < 2E-16 | 177.45 | 4.28 | 41.45 | < 2E-16 |
| Age | -0.81 | 0.29 | -2.78 | **0.005** | -1.59 | 0.32 | -5.03 | **5.35E-07** |
| Cohort (NORMENT) | 19.82 | 6.25 | 3.17 | **0.002** | 49.07 | 6.82 | 7.20 | **8.08E-13** |
| Age-by-cohort | 1.71 | 0.68 | 2.50 | **0.012** | 2.21 | 0.75 | 2.96 | **0.003** |

| sbTIV |  |  |  |  |
| --- | --- | --- | --- | --- |
|  | Estimate | Std. error | t-value | p-value |
| Intercept | 1,469.49 | 2.93 | 501.11 | < 2E-16 |
| Sex (M) | 155.68 | 4.06 | 38.35 | < 2E-16 |
| Age | -1.25 | 0.30 | -4.16 | **3.27E-05** |
| Cohort (NORMENT) | 9.31 | 6.47 | 1.44 | 0.150 |
| Age-by-cohort | 1.63 | 0.71 | 2.31 | **0.021** |

**Supplementary Table 11.** Fixed factors for the main longitudinal model for each ICV estimation method. Significant p-values are marked in bold type.

| eTIV |  |  |  |  |  | FSL |  |  |  |  |
| --- | --- | --- | --- | --- | --- | --- | --- | --- | --- | --- |
|  | Estimate | Std. error | DOF | t-value | p-value | Estimate | Std. error | DOF | t-value | p-value |
| Intercept | 1,396.96 | 44.60 | 2,333.62 | 31.33 | **< 2E-16** | 1,315.04 | 39.60 | 2,333.87 | 33.21 | **< 2E-16** |
| Sex (M) | 162.43 | 4.80 | 2,332.93 | 33.82 | **< 2E-16** | 164.47 | 4.26 | 2,332.88 | 38.57 | **< 2E-16** |
| Age | -1.65 | 0.34 | 2,350.40 | -4.92 | **9.40E-07** | -0.76 | 0.30 | 2,342.60 | -2.54 | **0.011** |
| Age<sup>2</sup> | -0.02 | 0.02 | 3,328.86 | -1.02 | 0.306 | -0.03 | 0.02 | 3,929.04 | -1.64 | 0.101 |
| Interscan interval | 4.04 | 19.58 | 2,332.07 | 0.21 | 0.836 | 3.15 | 17.38 | 2,331.65 | 0.18 | 0.856 |
| Scanner | -10.14 | 5.01 | 2,332.13 | -2.02 | **0.043** | -7.52 | 4.45 | 2,331.83 | -1.69 | 0.091 |
| Timepoint (Follow-up) | -10.91 | 1.01 | 4,602.60 | -10.81 | **< 2E-16** | -9.55 | 1.04 | 4,549.38 | -9.21 | **< 2E-16** |

| SPM12 |  |  |  |  |  | CAT12 |  |  |  |  |
| --- | --- | --- | --- | --- | --- | --- | --- | --- | --- | --- |
|  | Estimate | Std. error | DOF | t-value | p-value | Estimate | Std. error | DOF | t-value | p-value |
| Intercept | 1,382.94 | 38.22 | 2,333.55 | 36.19 | **< 2E-16** | 1,381.49 | 41.62 | 2,333.74 | 33.19 | **< 2E-16** |
| Sex (M) | 186.79 | 4.12 | 2,332.93 | 45.38 | **< 2E-16** | 176.77 | 4.48 | 2,333.03 | 39.44 | **< 2E-16** |
| Age | -0.88 | 0.29 | 2,349.94 | -3.06 | **0.002** | -1.63 | 0.31 | 2,347.14 | -5.20 | **2.13E-07** |
| Age<sup>2</sup> | -0.03 | 0.02 | 3,352.79 | -1.82 | 0.069 | -0.02 | 0.02 | 3,530.13 | -0.82 | 0.415 |
| Interscan interval | -1.14 | 16.78 | 2,331.97 | -0.07 | 0.946 | 7.53 | 18.27 | 2,331.96 | 0.41 | 0.680 |
| Scanner | -12.08 | 4.29 | 2,332.12 | -2.82 | **0.005** | -9.97 | 4.67 | 2,332.14 | -2.13 | **0.033** |
| Timepoint (Follow-up) | -10.75 | 0.87 | 4,615.64 | -12.36 | **< 2E-16** | -13.85 | 0.99 | 4,665.81 | -14.00 | **< 2E-16** |

| sbTIV |  |  |  |  |  |
| --- | --- | --- | --- | --- | --- |
|  | Estimate | Std. error | DOF | t-value | p-value |
| Intercept | 1,487.40 | 39.73 | 2,333.59 | 37.44 | **< 2E-16** |
| Sex (M) | 153.49 | 4.28 | 2,332.93 | 35.87 | **< 2E-16** |
| Age | -1.45 | 0.30 | 2,349.83 | -4.84 | **1.35E-06** |
| Age<sup>2</sup> | -0.07 | 0.02 | 3,358.51 | -4.31 | **1.71E-05** |
| Interscan interval | -4.62 | 17.44 | 2,332.00 | -0.27 | 0.791 |
| Scanner | -11.76 | 4.46 | 2,332.11 | -2.64 | **0.008** |
| Timepoint (Follow-up) | -10.09 | 0.91 | 4,618.53 | -11.14 | **< 2E-16** |

**Supplementary Figure 2.**
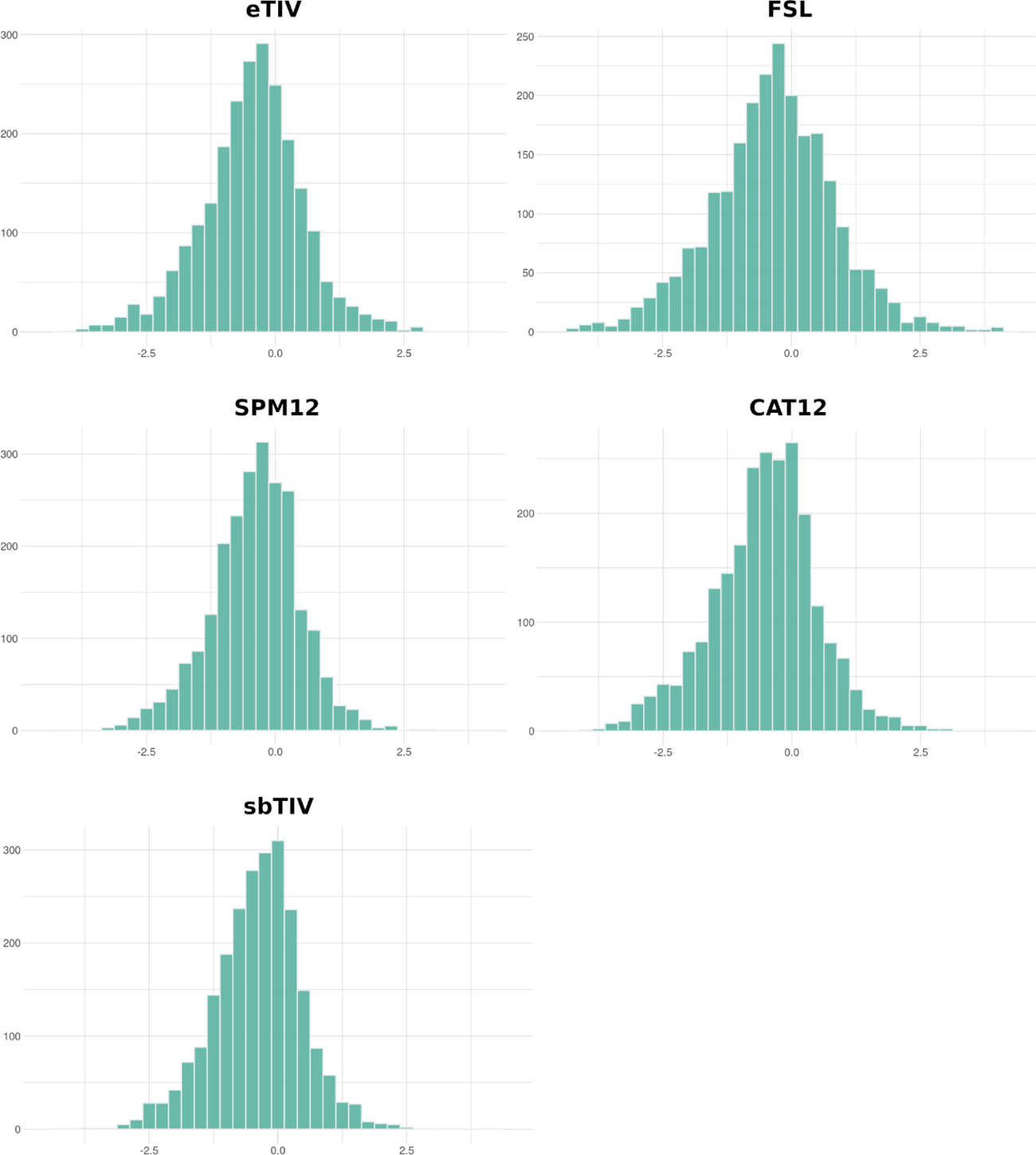
Histograms depicting the uncorrected annual percentage ICV differences between baseline and follow-up for each ICV estimation method.

